# Genetic and environmental influences on the evolution of virulence in the HIV-associated opportunistic human fungal pathogen *Cryptococcus neoformans*

**DOI:** 10.1101/2021.10.15.464490

**Authors:** Yiwu Yu, Yuanyuan Wang, Linghua Li, Xiaoqing Chen, Xinhua Huang, Huaping Liang, Tong Jiang, Guojian Liao, Min Chen, Liping Zhu, Muyuan Li, Tao Zhou, Qinyu Tang, Jingjun Zhao, Changbin Chen

**Affiliations:** Department of Dermatology, Tongji Hospital, Tongji University School of Medicine, Shanghai 200065, China; The Center for Microbes, Development and Health, Key Laboratory of Molecular Virology and Immunology, Institute Pasteur of Shanghai, Chinese Academy of Sciences, Shanghai 200031, China; Institute of Infectious Disease, Guangzhou No.8 People’s Hospital, Guangzhou 510060, Guangdong, China; Gusu School, Nanjing Medical University, Suzhou Municipal Hospital, The Affiliated Suzhou Hospital of Nanjing Medical University, Suzhou 215002, Jiangsu, China; State Key Laboratory of Trauma, Burns and Combined Injury, Department of Wound Infection and Drug, Army Medical Center (Daping Hospital), Army Medical University, Chongqing 400042, China; Nanjing Advanced Academy of Life and Health, Nanjing 211135, Jiangsu, China; University of Chinese Academy of Sciences, Beijing 100049, China; College of Pharmaceutical Sciences, Southwest University, Chongqing 400715, China; Department of Dermatology, Shanghai Key Laboratory of Medical Mycology, Changzheng Hospital, Second Military Medical University, Shanghai, China; Department of Infectious Diseases, Huashan Hospital, Fudan University, Shanghai 200040, China

**Keywords:** *Cryptococcus neoformans*, Virulence, Genotype-phenotype correlation, Environmental factors, Genetic variants

## Abstract

The fungus *Cryptococcus neoformans* is considered the leading cause of death in immunocompromised patients. Despite numerous investigations concerning its molecular epidemiology, there are only a few studies addressing the impacts of varying factors on genotype-phenotype correlations. It remains largely unknown whether genetic and environmental variabilities among isolates from different sources may have dramatic consequences on virulence. In this study, we analyzed 105 Chinese *C. neoformans* isolates, including 54 from HIV-infected patients, 44 from HIV-uninfected individuals and 7 from a natural environment, to investigate factors influencing the outcome of *C. neoformans* infection. MLST analysis clearly identified sequence type (ST) 5 as the prevalent sequence type in all clinical isolates and interestingly, genotypic diversities were observed in isolates from both HIV-uninfected individual and natural environment but not those from HIV-infected patients. Moreover, we found that compared to those from HIV-infected patients, the isolates from HIV-uninfected individuals exhibited enhanced virulence-associated traits including significantly elevated capsule production and melanin formation, increases in survival in human cerebrospinal fluid (CSF), less effective uptake by host phagocytes, and higher mortality in a mouse model of cryptococcosis. Consistently, pathogenic phenotypes were associated with CD4 counts of patients, implying environmental impact on within-host *C. neoformans* virulence. Importantly, a large-scale whole-genome sequencing analysis revealed that genomic variations within genes related to specific functions may act as a vital driving force of host intrinsic virulence evolution. Taken together, our results support a strong genotype-phenotype correlation suggesting that the pathogenic evolution of *C. neoformans* could be heavily affected by both genetic and environmental factors.

## Introduction

Cryptococcosis refers to a major invasive fungal infection caused by the encapsulated yeast species of the genus *Cryptococcus*, particularly *Cryptococcus neoformans* and *Cryptococcus gattii*. *C. neoformans* contains two varietal forms, *C. neoformans var. grubii* and *C. neoformans* var*. neoformans* [1, 2], and is distributed worldwide. In comparison, *C. gattii* was originally thought to be restricted only in tropical and subtropical districts but now this species was recognized in expanded temperate regions due to an outbreak of cryptococcosis on Vancouver Island, Canada [3]. The prevalence of cryptococcosis has increased in the past decade, and nearly a million cases of cryptococcal meningitis are diagnosed annually around the world, mainly in immunocompromised patients due to Human Immunodeficiency Virus (HIV) infection, organ transplantation, cytotoxic chemotherapy, and corticosteroid use [4, 5]. Clinical isolates of the *C. neoformans* and *C. gattii* species complexes are responsible globally for 15 to 30% of deaths in HIV/AIDS patients due to cryptococcal meningitis [6]. Meanwhile, *C. neoformans* accounts for the most common cause of meningitis in HIV adults in sub-Saharan Africa [6].

Phylogenetic analyses, as well as a number of genotyping studies, are being conducted using clinical and environmental *C. neoformans* isolates collected all over the world, in order to lay the basis for a comprehensive picture of the global genetic structure of this fungus [7, 8], and significant genetic diversities are observed in the *C. neoformans* species complex. All clinical *Cryptococcus* strains were initially treated as a single species with a conserved name *C. neoformans* [9]. It was later classified into four serotypes (A, B, C and D) based on Cryptococcal antigenic heterogeneity [10, 11]. Revised taxonomy further categorized the serotype B and C isolates to *C. gattii* [12], while *C. neoformans* encompassed three major serotypes including *C. neoformans* var*. grubii* (serotype A), *C. neoformans* var*. neoformans* (serotype D) and the hybrid serotype AD [1]. Moreover, the *C. neoformans* species complex consists of two evolutionary divergent species, *C. neoformans* and *C. deneoformans*, as well as their associative hybrids (*C. neoformans* x *C. deneoformans*). Of the two lineages, *C. neoformans* exhibited a worldwide distribution causing among 95% of cryptococcal infections and >99% in AIDS individuals, whereas *C. deneoformans* is more frequent in Europe and less virulent [13–15]. Due to a rapid development of molecular biology techniques including PCR fingerprinting, amplified fragment length polymorphism (AFLP) analysis, multilocus sequencing MLST and whole-genome sequencing [16–19], the relatedness of isolates at a molecular level has revolutionized our ability to differentiate among molecular types of the genus *Cryptococcus*. It is now well appreciated that *C. neoformans* can be divided into at least five major molecular types (AFLP1/VNI, AFLP1A/VNB/VNII, AFLP1B/VNII, AFLP3/VNIII and AFLP2/VNIV) [2, 18, 20]. Among them, *C. neoformans var. grubii* (serotype A/VNI) was found to naturally reside on avian excreta and trees, have a worldwide distribution, and contribute to over 80% of cryptococcosis [21]. Similar to the VNI clade, the VNII clade is also globally distributed. However, *C. neoformans var. grubii* (serotype A/VNB) was primarily found in sub-Saharan Africa and South America [22, 23]. Moreover, *C. neoformans var. neoformans* (serotype D/VNIV) was mostly found in Western Europe and South America and the *C. neoformans* hybrid (serotype AD/VNIII) showed a higher prevalence in the Mediterranean area of Europe [2, 24]. Collectively, these observations strongly suggest that molecular types of the *C. neoformans* isolates not only differ in their serological, epidemiological and ecological characteristics, but also exhibit diverse features on clinical presentations, antifungal susceptibility and therapeutic outcomes [16, 25, 26].

Recently, a growing body of evidence highlights the impact of *C. neoformans* genotype on the host disease outcome. For example, an epidemiological study by Wiesner *et al*. found that genotypes of *C. neoformans* isolates from HIV/AIDS patients in Uganda could be grouped into three distinct clonal clusters within the VNI clade and among them, both ST93 and ST77 isolates showed the highest mortality risk [27]. Comparatively, a similar study using clinical isolates from South African HIV/AIDS patients identified that ST32 isolates in the VNB clade exhibit worse patient outcome [28] and moreover, genotypic analysis of clinical *C. neoformans* isolates from Brazil revealed that ST93 in VNI clade is the most prevalent sequence type in HIV-infected patients [23]. However, a strikingly different link between isolate genotype and disease outcome was found in Asian countries, where *C. neoformans* infections are often observed in immunocompetent individuals [29] and ST5 represents the major sequence type in *C. neoformans* isolates from East Asian countries including China, Japan, Vietnam and South Korea [30, 31]. For example, a study using 136 Vietnamese clinical isolates of *C. neoformans var. grubii* revealed that ST5 isolates are responsible for 82% of infections in HIV uninfected patients, compared to only 35% cases in HIV infected patients [31]. In addition, MLST analysis of Chinese clinical *C. neoformans* isolates by different research groups also identified ST5 as the predominant sequence type [32]. Of course, notable exceptions exist. For instance, studies have indicated that ST4 and ST6 are the major MLST types in *C. neoformans var. grubii* isolates of Thailand whereas ST93 turns to be the dominant isolates from India and Indonesia [30, 33]. Interestingly, detailed information regarding genetic diversity of the isolates suggests that those from Thailand posit an evolutionary origin in African and the strains from China may have the same African origin but were expanded more flexibly and globally [5, 33]. Taken together, these studies highlight the presence of global genetic diversities in *C. neoformans* isolates and argue the correlation between pathogen genotype and patient phenotype. Of course, it is important to note that the patho-phenotypic variations of *C. neoformans* isolates cannot be fully explained by genotypic diversity, and other factors should also be considered.

Factors determining disease prevalence and species specificity are relatively unknown, but are speculated to be associated with the host immune response, genetic varieties, and virulence factors. For example, the type 1 helper T-cell (Th_1_) response could stimulate classical activation of macrophages and eliminate internalized cryptococcal cells, however, the type 2 helper T-cell (Th_2_) response was found to promote the disseminated, uncontrolled cryptococcal infection [34, 35], suggesting that patients exhibiting different immune responses to cryptococcal infections could yield varied clinical outcomes. Moreover, the distribution and prevalence of the molecular types appear to be highly relevant to geographical locations, the size of samples and host characteristics [1, 2]. For example, previous studies suggested that *C. neoformans* serotype A is one of the most common varieties and account for the majority of cryptococcal infections in Asia, especially in HIV-AIDS patients [34, 35]. In addition, studies showed that a list of virulence factors including the presence and size of the polysaccharide capsule, melanin production by laccase, cell size variation, growth at 37 °C and secretion of enzymes such as phospholipase, proteinase and urease, sphingolipid utilization, contribute to *C. neoformans* pathogenicity [36, 37]. However, it remains unclear whether these factors were correlated, and if so, how to affect the evolution of *C. neoformans* pathogenicity?

In an attempt to interpret the molecular epidemiology of *Cryptococcus* species, the strain differences in genotype and phenotype, as well as the impacts of genetic and environmental correlations on the evolution of fungal virulence, we analyzed a collection of 105 clinical and environmental isolates of *C. neoformans* in China, including those isolated from HIV-infected patients and HIV-uninfected individuals. The genotype of each isolate was determined by MLST and the evaluation of pathogenic phenotypes was performed *in vitr*o and *in vivo*. Moreover, the genotype-phenotype correlations were further assessed by genetic variations through the genome-wide linkage and association analyses. Our results compare the impacts of genetic and environmental factors on affecting the correlation between genotype and phenotype of *C. neoformans* isolates, and provide *in vitro* and *in vivo* data to support the influence of genetic and environmental changes on genotypic and pathogenic variations.

## Materials and methods

### Ethics statement

All of animal experiments were performed in compliance with the Regulations for the Care and Use of Laboratory Animals issued by the Ministry of Science and Technology of the People’s Republic of China, which enforces the ethical use of animals. The protocol was approved by IACUC at the Institut Pasteur of Shanghai, Chinese Academy of Sciences (Permit Number: 160651A).

### Strains

A total of 105 isolates of *C. neoformans* strains were assayed in this work, including 44 from the HIV-uninfected patients (HIV-u group), 54 from the HIV-infected patients (HIV-i group) and 7 from the nature (Env group). Clinical and laboratory records of all patients were obtained from Guangzhou No.8 People’s Hospital, Huashan Hospital, Changzheng Hospital and Southwest University. The data collected for analysis included age, gender, initial symptoms, HIV infection status and CD4^+^ T cell count (at the time of diagnosis). The detailed information about each of the samples is presented in **S1 Table**. Among these, 98 strains were isolated from cerebrospinal fluid (CSF) samples (n = 90) and blood cultures (n = 8). *C. neoformans* clinical isolates were single colony purified on YPD medium and then maintained as glycerol stocks at −70°C for long-term storage. A detailed information about each sample is listed in **S2 Table**. Each strain was streaked and grown as a single colony on yeast peptone dextrose (YPD) medium prior to use.

A list of reference strains was included in this study, including international strains used for phylogenetic analysis (H99 in USA, WM148 and WM626 in Australia, ST93 in Brazil) and standard strains for *in vitro* and *in vivo* assays (JEC20 and H99). All strains were maintained on yeast peptone dextrose (YPD) medium prior to use.

### DNA extraction

Isolates were grown on Sabouraud dextrose agar slants for 48 h. Single colonies were isolated, re-inoculated in10ml of liquid medium, and grown at 30°C for 24 h. The cells were collected by centrifugation and genomic DNA was extracted using the cetyltrimethylammonium bromide (CTAB) DNA isolation method as previously described [38].

### Multilocus Sequence Typing (MLST)

Multilocus sequence type analysis was carried out using the seven ISHAM consensus loci (*CAP59*, *GPD1*, *IGS1*, *LAC1*, *PLB1*, *SOD1*, *URA5*), following a standard procedure [18, 22]. All primers used in this study were described previously [27] and listed in **S6 Table**.

### Phylogenetic analysis

The sequences were aligned with the computer program CLUSTAL W and a phylogenetic tree was drawn based on the neighbor-joining (NJ) model. The evolutionary distances were computed based on the *p*-distance, and all gaps were eliminated. A bootstrap analysis was performed with 1,000 replicates [39].

### Assays for melanization and capsule formation

Melanin production was measured by comparing pigmentation of strains grown on L-DOPA media to reference strains with strong (H99) and weak (JEC20) melanization, using a method described previously [40]. Relative melanization scores of zero (equal to JEC20), one to four (between JEC20 and H99) and five (more than or equal to H99) were assigned to the strains based on comparison with the reference strains grown on the same plate.

The capsule induction assay was performed using a method described previously [41], with some modifications. In brief, stationary-phase fungal cultures were washed and resuspended in PBS. Cells were diluted 1/100 in capsule induction medium [10% Sabouraud dextrose medium in 50mM MOPS buffer (pH 7.3)] and incubated at 30 °C and 180rpm for 48h. The size of capsule was measured by staining the cells with India ink and imaging at a magnification of ×63 under a light microscope. As described by Fernandes *et al*. [42], the diameters of the whole cell (yeast cell + capsule) and the cell body (the cell wall only) were each measured using Image J software, and the mean of the values was calculated for 10-20 cells for each isolate.

The identification of mating types was carried out in a classical PCR analysis using the mating type and serotype specific primers [43]. All primers are listed in **S6 Table**.

### Infection of macrophages with *Cryptococcus*

Phagocytosis assays were performed with the murine macrophage-like cell line J774, using a method previously described [44, 45]. Briefly, macrophage cells in DMEM supplemented with 10% heat-inactivated fetal bovine serum (FBS), 1 mM L-glutamine, and 1% penicillin-streptomycin (Sigma-Aldrich) were plated into each well of a 24-well tissue culture plate for 24 hours at 37°C with 5% CO_2_. Macrophage cells (1.5 × 10^5^) were incubated in serum-free DMEM medium for 2 hours, activated with 15μg/ml phorbol myristate acetate (PMA) (Sigma-Aldrich) for 30-60 minutes, and then co-incubated with *C. neoformans* yeast cells opsonized by a monoclonal antibody (C66441M, purchased from Meridian Life Science, Inc) for 2 hours at 37°C with 5% CO_2_ (MOI=1:10). Extracellular yeast cells were removed by extensive washes with prewarmed PBS. The extent of *Cryptococcus* phagocytosis was calculated as the number of cryptococci internalized by macrophages 2 hours after infection. Results were expressed as the mean of 3 to 4 experimental repeats.

### CSF survival assay

Human cerebrospinal fluid (CSF) was pooled from Shanghai Changzheng Hospital patients receiving serial therapeutic lumbar punctures, collected anonymously from populations of at least 15 patients. The human CSF was fully examined for parameters, including white cell count, protein and glucose levels, within the normal ranges and for the absence of antifungal drugs. Clinical isolates of *C. neoformans* were replicated in 48-well plates in Sabouraud dextrose broth (SDB) and incubated at 37°C for 3-4 days until saturation. Cultures were then diluted, inoculated into CSF at a concentration of 1~2 × 10^6^ cells/ml, and incubated at 37°C for 96 hours. As described previously [45], aliquots were collected at different time points (0,12, 24, 36,72 and 96 hours after inoculation) and plated on Sabouraud dextrose agar (SDA) media for CFU counts. The survival slope was determined as the mean rate of increase or decrease in cryptococcal counts after CSF treatment, by averaging the slope of the linear regression of log_10_ CFU/ml over time for each strain.

### Virulence studies

A well characterized murine inhalation model of cryptococcosis was used [46]. Briefly, three isolates were randomly picked from each of the three groups and individually grown overnight in liquid YPD cultures at 30 °C. All strains tested are ST5, except for one environmental strain (Env #103; ST39). Cells were counted using a hemocytometer and the yeast suspension with a final concentration of 1 × 10^7^ cells/ml was prepared. 6-8 week-old female C57BL/6 mice were anesthetized by intraperitoneal injection of ketamine (75 mg/kg) and medetomidine (0.5-1.0 mg/kg) and a 50 μL volume of the yeast suspension (1 × 10^5^ cells) was intranasally injected. 8 mice were infected per inoculum. Mice were monitored several times a week until the observance of disease symptoms (weight loss, ruffled fur, shallow breathing, abnormal gait and lethargy) and then monitored daily. Mice were sacrificed by CO_2_ inhalation followed by cervical dislocation when signs of severe morbidity, including significant weight loss, abnormal gait, hunched posture and swelling of the cranium, were clearly displayed. A Kaplan-Meier method was employed to analyze survival curves using GraphPad Prism 6.0 software. Differences in the median survival among species were determined by performing the log-rank test [38, 47].

### Whole Genome Sequencing

28 C. neoformans strains isolated from HIV-infected patients and HIV-uninfected patients were subjected to whole-genome sequencing. In order to minimize the genotype impact on the sequencing, all 28 strain were selected ST5. Next generation sequencing library preparations were constructed following the manufacturer’s protocol (NEBNext^®^ Ultra™ DNA Library Prep Kit for Illumina^®^). For each sample, 1 μg genomic DNA was randomly fragmented to <500 bp by sonication (Covaris S220) and DNA fragments were treated with End Prep Enzyme Mix for end repairing, 5’ Phosphorylation and dA-tailing in one reaction, followed by a T-A ligation to add adaptors to both ends. Size selection of Adaptor-ligated DNA was then performed using AxyPrep Mag PCR Clean-up (Axygen) and fragments of ~410 bp (with the approximate insert size of 350 bp) were recovered. Each sample was then amplified by PCR for 8 cycles using P5 and P7 primers, with both primers carrying sequences which can anneal with flowcell to perform bridge PCR and P7 primer carrying a six-base index allowing for multiplexing. The PCR products were cleaned up using AxyPrep Mag PCR Clean-up (Axygen), validated using an Agilent 2100 Bioanalyzer (Agilent Technologies, PaloAlto, CA, USA), and quantified by Qubit2.0 Fluorometer (Invitrogen, Carlsbad, CA, USA). The whole genome of each strain was sequenced using Illumina NovaSeq6000 PE150 at the Beijing Novogene Bioinformatics Technology Co., Ltd.

### Data analysis

The raw data was removed the sequences of adaptors, polymerase chain reaction (PCR) primers, content of N bases more than 10% and bases of quality lower than 20 by using Cutadapt (V1.9.1). Then the clean data was mapped to the *C. neoformans* H99 reference genome FungiDB (http://fungidb.org/fungidb/) by using BWA (V0.7.17) and mapping results were processed by Picard (V1.119) to remove duplication. SNPs/InDels were called by the GATK Unified Genotyper (V3.8.1) and Annotated by Annovar (V21 Apr 2018). GO-term analysis of processes enriched among specific gene sets of HIV-i and HIV-u group was performed using the topGO (V2.34.0).

### Statistics

Data were presented as mean ± standard error of the mean (SEM). Statistical analysis was performed by one-way analysis of variance (ANOVA) in GraphPad Prism 6.0 software (San Diego, CA). The following *p*-values were considered: * *p* < 0.05; ** *p* <0.01; *** *p* < 0.001; **** *p* < 0.0001.

### Data availability

The authors declare that the data supporting the findings of this study are available within the article and its supplementary information files, or are available upon request to the corresponding authors. Whole-genome sequences are available at NCBI under BioProject ID: PRJNA680424.

## Results

### Characterization of clinical presentation and outcome

In order to study the correlations amongst environmental conditions, immune inflammatory syndrome and other clinical outcomes, we collected a total of 98 clinical isolates of *C. neoformans* that were recovered from cerebrospinal fluid (CSF) samples (n=90) and blood cultures (n=8) of 98 patients hospitalized at different locations of China. Among them, 54 strains are from HIV-infected patients and 44 are from HIV-uninfected individuals. Clinical laboratory data was investigated from 95 patients (three patients’ data was somehow missed). Both **Table 1** and **S1 Table** showed the detailed information about the influence of common symptoms on clinical outcome that could be used to Fig out the relationships between isolate origin and clinical consequences. The majority of patients were male (74; 77.9%), the age ranged from 21 to 68 years and the mean age was 41 ± 11 years.

**Table 1.**
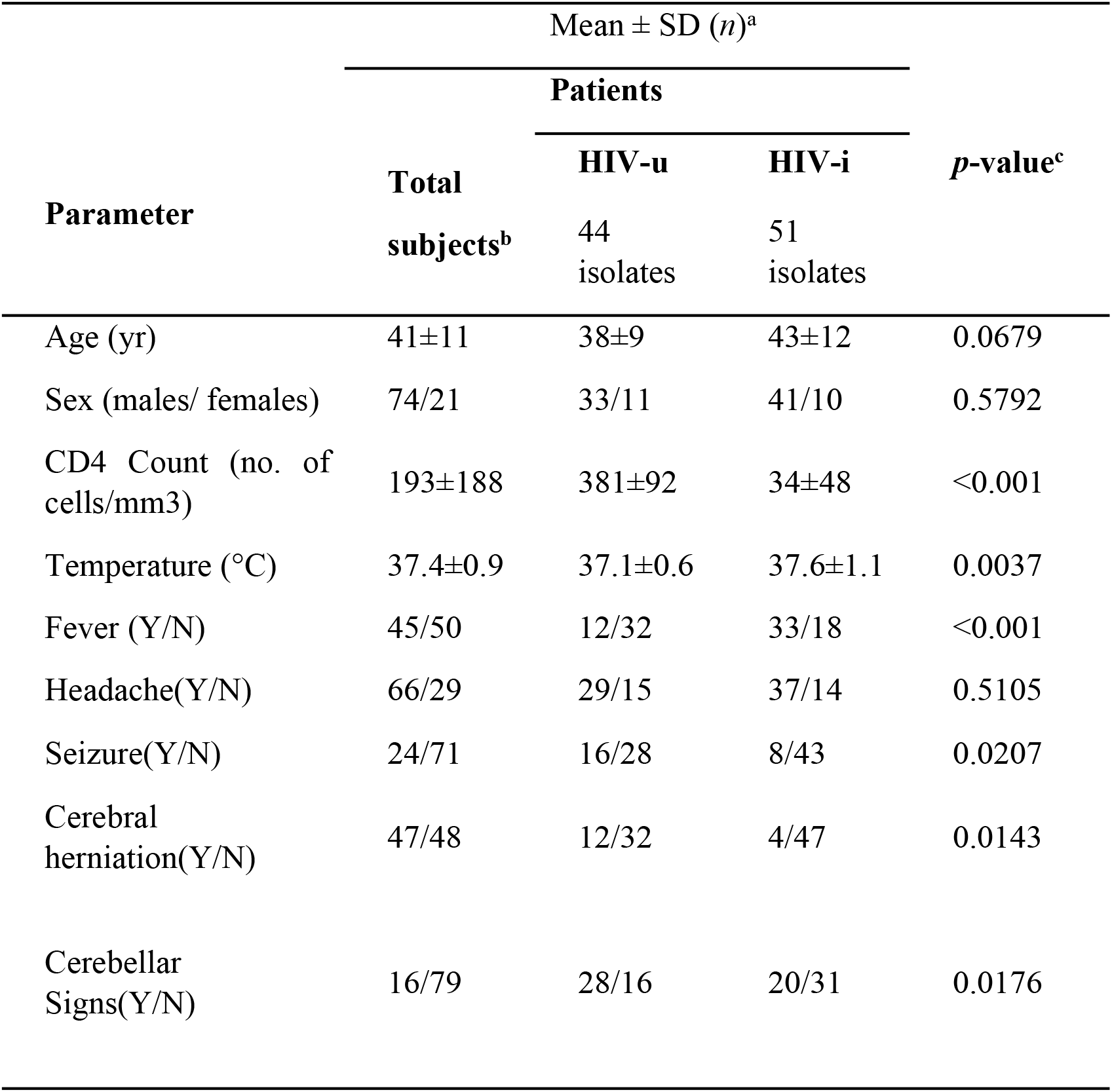
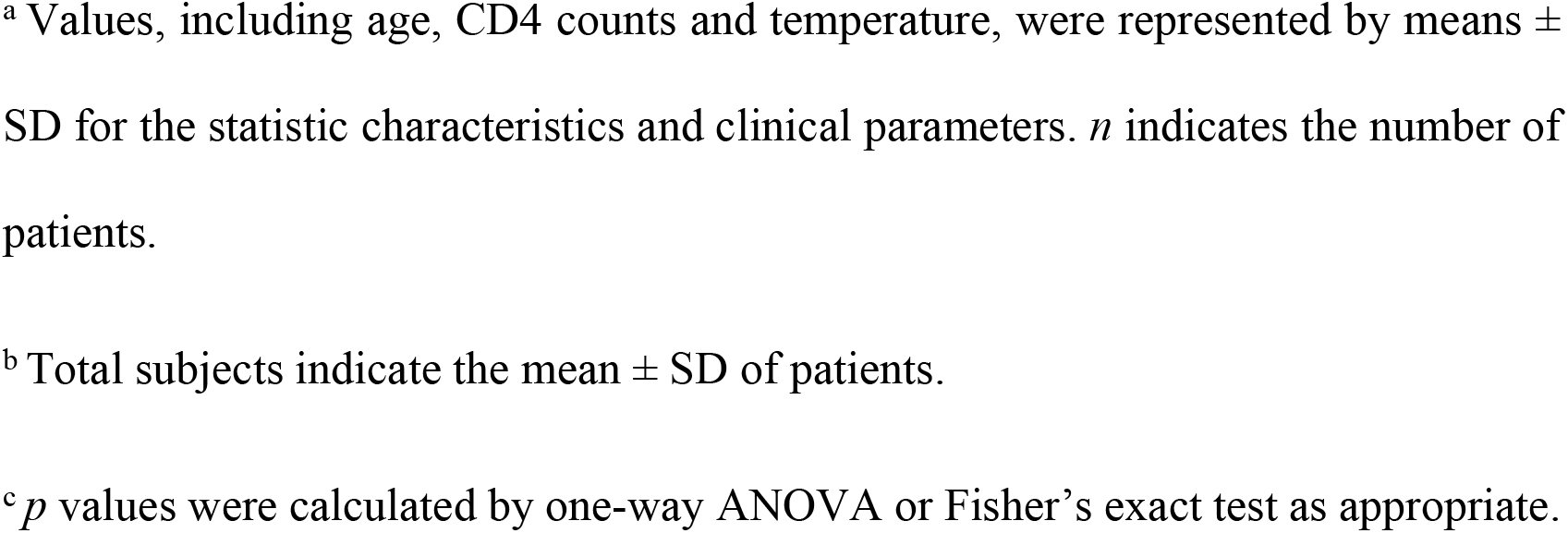
Comparison of clinical data for HIV-uninfected and -infected groups

| Parameter | Mean ± SD (*n*)<sup>a</sup> | | | *p*-value<sup>c</sup> |
| --- | --- | --- | --- | --- |
|  | Total<br>subjects<sup>b</sup> | Patients |  |  |
|  |  | HIV-u<br>44<br>isolates | HIV-i<br>51<br>isolates |  |
| Age (yr) | 41±11 | 38±9 | 43±12 | 0.0679 |
| Sex (males/ females) | 74/21 | 33/11 | 41/10 | 0.5792 |
| CD4 Count (no. of<br>cells/mm3) | 193±188 | 381±92 | 34±48 | <0.001 |
| Temperature (°C) | 37.4±0.9 | 37.1±0.6 | 37.6±1.1 | 0.0037 |
| Fever (Y/N) | 45/50 | 12/32 | 33/18 | <0.001 |
| Headache(Y/N) | 66/29 | 29/15 | 37/14 | 0.5105 |
| Seizure(Y/N) | 24/71 | 16/28 | 8/43 | 0.0207 |
| Cerebral<br>herniation(Y/N) | 47/48 | 12/32 | 4/47 | 0.0143 |
| Cerebellar<br>Signs(Y/N) | 16/79 | 28/16 | 20/31 | 0.0176 |

<sup>a</sup> Values, including age, CD4 counts and temperature, were represented by means ± SD for the statistic characteristics and clinical parameters. *n* indicates the number of patients.
<sup>b</sup> Total subjects indicate the mean ± SD of patients.
<sup>c</sup> *p* values were calculated by one-way ANOVA or Fisher's exact test as appropriate.

Among them, the most common presenting symptom was headache (70%), especially in HIV-infected patients (73%). Comparatively, the HIV-uninfected patients were less frequently found to have high fever but they had more severe symptoms such as seizure, cerebral herniation and cerebellar signs. Moreover, 39 out of 51 HIV-infected patients presented CD4^+^ T-cell counts below 50 /mm^3^. However, the mean ± SD of CD4^+^ T-cell counts from 44 HIV-uninfected individuals was 381 ± 92 mm^3^/ml. These results are consistent with a recent study that the HIV-associated cryptococcal meningitis normally occurs in patients with higher CD4 counts [48].

### MLST and phylogenetic analyses

The 98 clinical isolates, together with 7 environmental strains collected from pigeon excreta, were classified into three groups (HIV-u group: 44 strains from HIV-uninfected individuals; HIV-i group: 54 strains from HIV-infected patients; Env group: 7 environmental strains from the pigeon excreta). After colony purification, all 105 isolates were analyzed to determine their genotypes by Multilocus Sequence Typing (MLST), based on a formal study that described a consensus sequence-based epidemiological typing scheme, using seven housekeeping genes (*CAP59*, *GPD1*, *IGS1*, *PLB1*, *LAC1*, *SOD1*, *URA5*) [18]. For each strain, locus allele identifiers were used and combined to generate a sequence type (ST) representing the strain genotype [49]. MLST analysis of all 105 isolates demonstrated the presence of 9 sequence types (ST), including ST5 (n = 94; 89.5%), ST359 (n = 2; 1.9%), ST2 (n = 2; 1.9%), and ST39 (n = 2; 1.9%), ST360 (n=1; 0.9%), ST194 (n=1; 0.9%), ST31 (n=1; 0.9%), ST93 (n=1; 0.9%), ST195 (n=1; 0.9%) (**Fig 1A**). To our surprise, we found in **S2 Table** that all isolates from HIV-i group were restricted to a single ST (ST5). In comparison, STs identified in isolates from HIV-u and Env groups were more diverse, as we observed that strains from HIV-u group contain 8 STs, including ST5 (n=36), ST359 (n=2), ST2 (n=1), ST360 (n=1), ST194 (n=1), ST31 (n=1), ST93 (n=1) and ST195 (n=1), and strains from the Env group also harbor 3 STs, including ST5 (n=4), ST39 (n=2) and ST2 (n=1).

**Fig 1.**
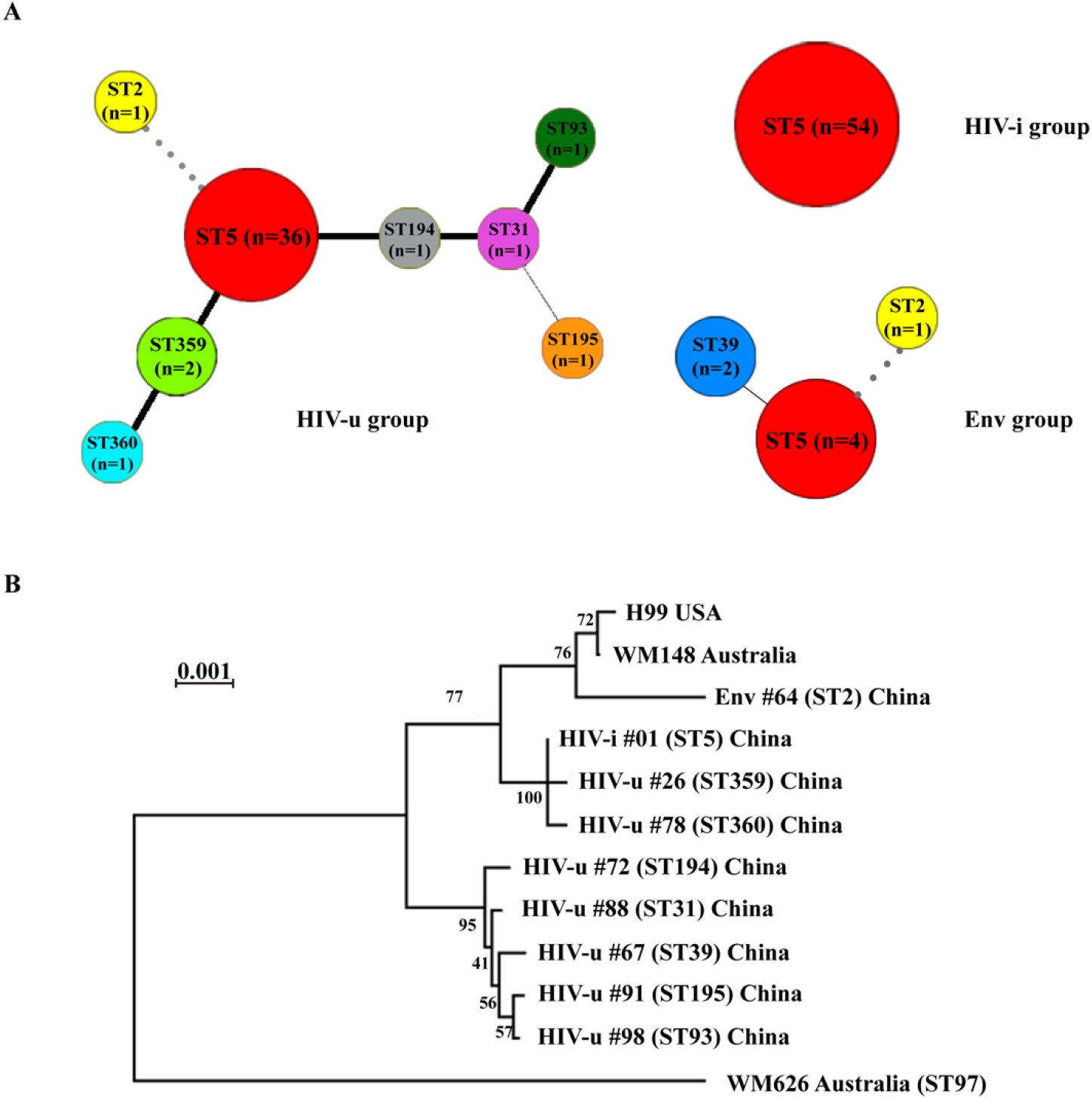
Genetic and evolutionary relationships among *C. neoformans* strains derived from different sources. **(A)** Hypothesis of the evolution of the 105 C. neoformans strains, based on phylogenetic relationships described by sequence analysis of the MLST scheme. Representative STs labelled with different colors. The number of isolates sharing the same ST is listed in each node, whereas the lines between STs indicate inferred phylogenetic relationships and are denoted by bold black, plain black, or grey line, depending on the number of allelic mismatches between profiles (bold black: < 3; Plain black: 3-4; Grey: > 4). **(B)** Phylogeny of selected *C. neoformans* strains based on DNA sequences from eight housekeeping gene loci. The tree was constructed with the allelic frequencies using the neighbor-joining (NJ) model and 1,000 bootstrap replicates were performed. Numbers labeled with HIV-u, HIV-i or Env are strains from 105 Chinese isolates in this study; Three reference strains, one from the United States and the other two from Australia, are also included.

In addition, a phylogenetic analysis was performed among strains from different geographic locations (China, Australia, Brazil and USA), based on the concatenated sequences of MLST loci (**Fig 1B**). The phylogenetic tree revealed that ST5 was quite close to ST359 and ST360, while ST39 was far from other identified STs. Moreover, we concluded from this study that the major epidemic clone of *C. neoformans var. grubii* in China was ST5 regardless of their origins. Actually, similar patterns were observed in previous studies using strains isolated from HIV-uninfected individuals of China [7, 50].

As shown in **Fig 1A**, we found that the majority of the isolates in this collection were of one sequence type (ST5), which was 54 out of 54 in the HIV-i group, 36 out of 44 in the HIV-u group and 4 out of 7 in the Env group, prompting us to evaluate the potential relevance of the isolates to virulence according to their origins other than the STs.

### *In vitro* and *ex vivo* phenotyping

*C. neoformans* produces several important virulence factors, most notably the mating type, polysaccharide capsule, and melanin production. Studies have shown that mating type can influence virulence through cell type (*MAT*α or *MAT***a**) and the function of specific genes such as those related to MAP kinase pathway [51–53]. The mating types and serotypes of all isolates were determined by multiplex PCR using specific primers described previously [18]. Among the 105 strains evaluated, 104 were characterized as serotype A MATα (Aα) and only one strain was found to be serotype A MAT**a** (A**a**) (**S1 Fig**).

Melanin is another major virulence factors in *C. neoformans*. We carried out *in vitro* assays to evaluate the ability of all 105 isolates to produce melanin by patching on medium containing L-DOPA (L-3,4-dihydroxyphenylalanine) [54]. A scoring method, based on K-means clustering analysis, was employed to evaluate melanin production of all tested isolates. Two reference strains (JEC20 and H99) were used as experimental controls, given the fact that the strain JEC20 has been reported to produce almost no melanin whereas the clinical isolate H99 exhibits strong melanization [55]. As shown in **Fig 2 and S2 Fig**, we found that strains in HIV-u group produce significantly higher levels of melanin, as showed by dark pigments, when compared to those in HIV-i group (*p* < 0.0005 by Tukey adjusted t-test), implying that the clinical isolates from the two groups may have differences in pathogenicity.

**Fig 2.**
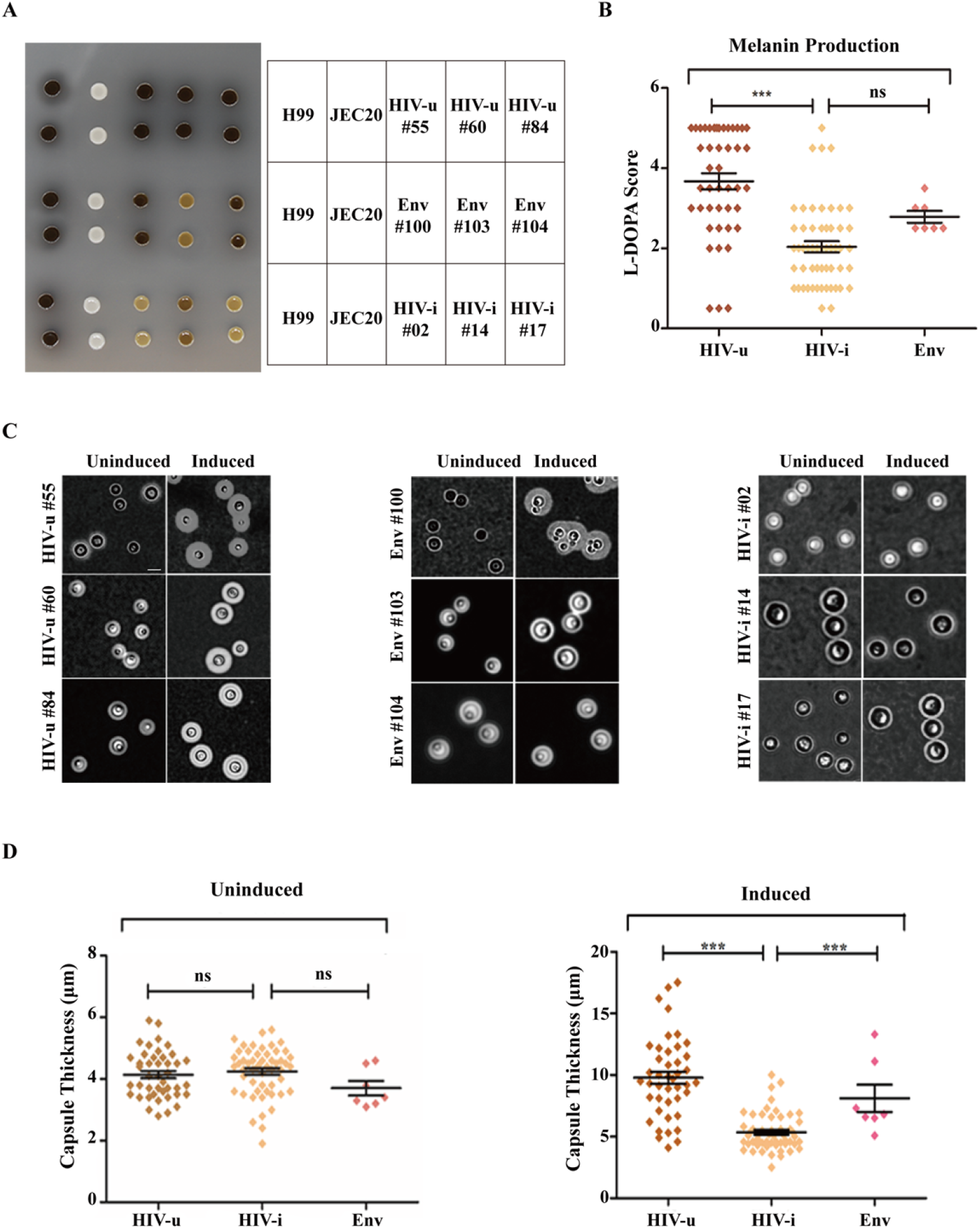
*In vitro* characterization of melanin biosynthesis and the formation of polysaccharide capsule in *Cryptococcus* isolates from each of the three groups. **(A)** Three representative isolates were randomly selected from each group and melanin phenotypes were verified by growth at 30 °C on plates containing L-DOPA. Pictures were taken after incubation for 3 days. *C. neoformans* strains H99 and JEC20 were used as positive and negative control, respectively. There were two replicates for each strain. **B** Statistical comparison of melanin biosynthesis in all 105 isolates of *C. neoformans*. The scores were calculated based on a K-Means cluster analysis. The vertical bars represent standard errors of the means in each group. ns *p* > 0.05, *** *p* < 0.001. **(C)** As in A, the same isolates were analyzed for capsule formation by India ink staining under capsule-inducing and non-inducing growth conditions. A single colony from each isolate was resuspended in PBS containing India ink, subjected to a short vortexing and examined under a light microscope at ×63 magnification. Scale bars, 10 μm. (D) Statistical comparison of capsule formation in all 105 isolates of *C. neoformans*. As described in Materials and Methods, stationary-phase fungal cultures were washed and resuspended in PBS, and diluted 1/100 in capsule induction medium (10% Sabouraud dextrose medium in MOPS buffered at pH 7.3). After incubation at 30 °C and 180rpm for 48h, suspensions of India ink were photographed and the capsule thickness of each isolate was measured. Each symbol represents the average capsule thickness of 10-20 cells. \**p* < 0.05, \*\*\**p* < 0.001, by unpaired two-tailed Student’s test.

This notion was further supported by assaying the polysaccharide capsule formation of each isolate. Capsule induction of each isolate was achieved under nutrient-limiting conditions (capsule induction medium; 10% Sabouraud dextrose medium in MOPS buffered at pH 7.3) and measured by staining with India ink, following a protocol described in Materials and Methods. The capsule size of each strain was determined by Adobe Photoshop software (Adobe Inc, USA) and the data was further analyzed by a K-means clustering algorithm. We observed that after induction, strains in HIV-u group generate much larger capsules than those in HIV-i group, although their sizes are almost indistinguishable under un-induced condition (**Fig 2 and S3 Fig**).

The ability of *C. neoformans* to survive in the cerebrospinal fluid (CSF) has been found to contribute to the cryptococcal virulence due to its presence in clinical specimens and production of life-threatening disease in the central nervous system (CNS). To further evaluate the pathogenic variabilities between cryptococcal strains from HIV-infected and uninfected individuals, we assayed the survival of all 98 clinical isolates in human CSF. When compared to those from the HIV-I group, isolates from the HIV-u group exhibited more resistant to killing by human CSF (**Fig 3A**). Moreover, we found a positive correlation between CSF survival and capsule size (*r =*0.6737, *p* <0.0001;**Fig 3B**). These data sustain the proposition that strains in HIV-u group appear to be more virulent than host in HIV-I group.

**Fig 3.**
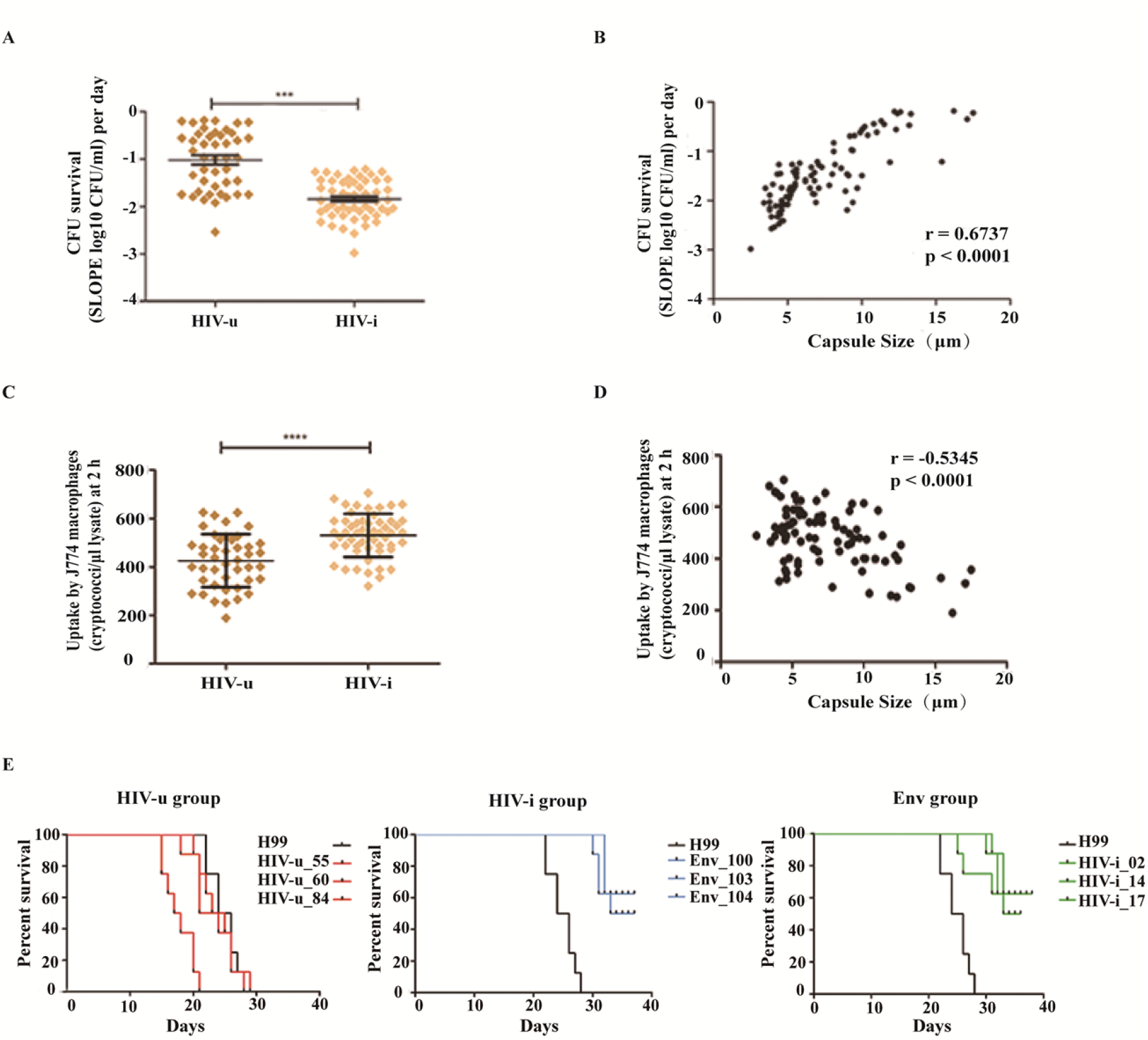
*Ex vivo* and *in vivo* assays investigating virulence-associated characteristics in *Cryptococcus* isolates from each of the three groups. **(A, B)** Cryptococcal survival in human CSF. Cells of each clinical isolate were inoculated into human CSF at a concentration of 1~2 × 10^6^ cells/ml and aliquots were collected at different time points (0,12, 24, 36,72 and 96 hours after inoculation) and plated on Sabouraud dextrose agar (SDA) media for CFU counts. The survival slope was determined as the mean rate of increase or decrease in cryptococcal counts after CSF treatment, by averaging the slope of the linear regression of log_10_ CFU/ml over time for each strain. A. Survival comparison of clinical *C. neoforman*s strains between HIV-u and HIV-i group after incubation with human CSF; B. Association of capsule size with *ex vivo* cryptococcal survival in human CSF. **(C, D)** Phagocytic uptake of clinical *C. neoforman*s strains by macrophage-like cell line J774. Macrophage cells (1.5 × 10^5^) were incubated in serum-free DMEM medium for 2 hours, activated with 15μg/ml phorbol myristate acetate (PMA) for 30-60 minutes, and then co-incubated with *C. neoformans* yeast cells pre-opsonized by a monoclonal antibody for 2 hours at 37°C with 5% CO_2_ (MOI=1:10). The extent of *Cryptococcus* phagocytosis was calculated as the number of cryptococci internalized by macrophages 2 hours after infection. Results were expressed as the mean of 3 to 4 experimental repeats. C. Phagocytic uptake comparison of clinical *C. neoforman*s strains between HIV-u and HIV-i group after incubation with J774; D. Association of capsule size with *ex vivo* phagocytic uptake. **(E)** Kaplan-Meier survival curves of mice infected with individual strains. Groups of female C57BL/6 mice (8 for each group) were inoculated intranasally with 1×10^5^ CFUs of the indicated strain and monitored for progression to severe morbidity. As in Fig 2, the same three representative isolates from each of the three groups were used. For comparison, the strain H99 was used as a control. (Left) HIV-u group; (Middle) HIV-i group; and (Right) Env group. Note: All *C. neoformans* strains, including the control strain H99 and all 9 tested isolates, were tested for pathogenicity in one experiment. \*\*\**p* < 0.001, by Log-rank test.

In addition, many studies have emphasized the role of fungal internalization by macrophages as a critical virulence factor in cryptococcal disease [56, 57]. Interestingly, when the macrophage uptake of *C. neoformans* cells was evaluated in the established murine macrophage-like cell line J774, we observed that isolates from the HIV-i group were phagocytosed at a higher rate than those from the HIV-u group (**Fig 3C**), and this trait could be explained by differences in capsule size, since phagocytic uptake and capsule size is inversely correlated in this set of isolates (*r* = – 0.5345, *p* <0.0001;**Fig 3D**). Actually, our results are consistent with previous reports that the capsule passively inhibits phagocytosis of *C. neoformans* by macrophages and the capsule-dependent anti-phagocytic activity represents a major virulence attribute [45, 58, 59].

Taken together, both *in vitro* and *ex vivo* phenotypic assays suggest that compared to those from the HIV-infected patients, the clinical isolates derived from the HIV-uninfected individuals exhibit significantly enhanced capsule production and melanin formation, higher increases in survival in CSF, and less effective uptake by host phagocytes, which represent key factors associated with *Cryptococcus* pathogenicity [60, 61].

### *In vivo* virulence analyses

Given the observed differences in capsule and melanin production between strains from HIV-uninfected and infected groups, we sought to ask whether strain origins may affect *C. neoformans* virulence. We conducted an *in vivo* virulence analysis using a mouse model of cryptococcosis [46]. Groups of 6-8-week female C57BL/6 mice (8 mice per group) were inoculated intranasally with strains from the three groups (we randomly picked three strains in each group), as well as the H99 strain serving as a control. All isolates tested are ST5, except for one environmental strain (Env #103; ST39). Mice were sacrificed by CO_2_ inhalation followed by cervical dislocation when signs of severe morbidity, including weight loss, abnormal gait, hunched posture and swelling of the cranium, were clearly observed. Consistent with a previous study [46], the control strain H99 caused a lethal infection by day 20. As expected, we observed that mice inoculated with the strains of HIV-u group showed similar or even worse signs of disease as the control strain. In contrast, mice inoculated with strains from both HIV-i and Env groups showed almost identical survival rates (**Fig 3E**), and the strains in these two groups are much less virulent than those in HIV-u group.

Thus, consistent with the results obtained from *in vitro* and *ex vivo* assays, our *in vivo* analysis also indicated that clinical isolates of *C. neoformans* from different hosts exhibit differentiation of virulence phenotypes, implying the importance of environmental factors on the evolution of virulence.

### Genetic variations

Studies have shown that microbial pathogens evolve a broad range of intrinsic and extrinsic strategies to acquire and modulate existing virulence traits in order to achieve successful colonization in the host [62]. In addition to the impact of environmental factors, pathogenicity of *C. neoformans* must be also influenced at the genetic level, since genetic variation in the genome is recognized as a fact of life in microbes, allowing pathogens to adapt to their chosen host and also to resist clearance by the host. Moreover, whole genome-resequencing technology has been used as a powerful tool to explore the genetic mechanism in fungi and fungal-host interaction [63]. Hence, we sequenced and assembled the whole genomes of 28 strains in the collection (ST5), 14 from HIV-u group and 14 from HIV-i group (the strains were chosen based on the patients’ parameters, including close age, sex and several other factors within the same group), and performed a comparative study to identify potential correlations between genetic variation and virulence, by comparing the genome difference of 28 isolates against the finished reference strain H99. Quality control analyses of our sequencing data were summarized in **S3 Table**. Overall, sequencing the 28 isolates yielded a total of 65 Gb containing 436 million raw reads, with an average of 15.6 million reads for each sample. The raw reads were further filtered by removing the low quality reads and duplicate reads, producing 52 Gb containing 349 million high quality reads. Approximate 94.8% of the clean reads could be mapped to the H99 reference genome, varied from 30.77 to 99.08% among different strains (**S4 Table**). These mapped sequences were used for further analyses.

**S5 Table** lists the number of mutations observed in the 28 clinical isolates comparing with the reference genome of H99. We identified a total of 65,106 variants, including 57,401 SNPs (single nucleotide polymorphism; 13,133 (22.9%) synonymous and 10,755 (18.7%) nonsynonymous) and 7,705 indels (insertions/deletions; 4,149 (53.8%) InDel-Ins and 3,556 (46.2%) InDel-Del), across the genomes of 14 HIV-u isolates. In comparison, our analysis across the genomes of 14 HIV-i isolates generated a total of 64,138 variants, with 56,655 SNPs (13,064 (23.1%) synonymous and 10,683 (18.9%) nonsynonymous) and 7,483 indels (4,058 (54.2%) InDel-Ins and 3,425 (45.8%) InDel-Del). The relationship between number of variants and gene was identified in **Fig 4A** and the inset panel was magnified to show genes with at least 20 variants for visualization purposes. The results showed that there was no significant difference in both groups. As shown in **Fig 4B**, the majority of genes in both groups harbor relatively few variants and there is no correlation between gene length and the number of variants, indicating that the relative frequencies of genetic variations in these clinical isolates are quite similar and appear to be independent of strain origins. In addition, we surprisingly found in **Fig 4C** that the genes harboring more than 80 variants were almost identical in all sequenced isolates of both groups, suggesting that differential virulence may not be generalized to regions encompassing extreme genomic instability, instead, may be attributed to genomic mutations of genes associated with specific functions.

**Fig 4.**
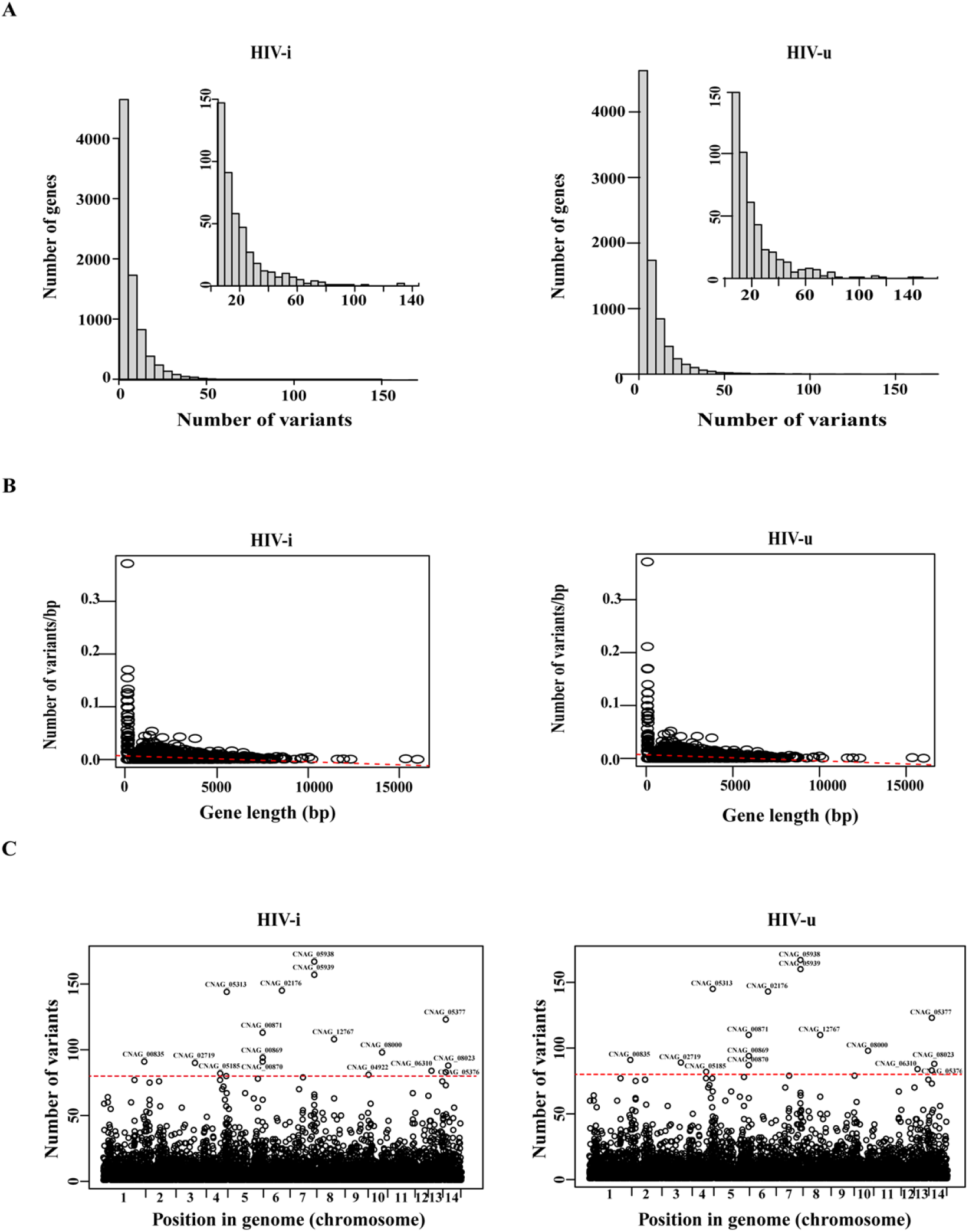
Summary of genomic variants identified from whole-genome sequencing of selected *C. neoformans* isolates. **(A)** A total of 28 clinical isolates, including 14 from the HIV-i group and another 14 from the HIV-u group, were sequenced. Shown is the number of variants that were identified in each gene locus. Genes with at least 20 variants were selected and presented in the magnified inset panel. **(B)** All sequenced isolates from both groups showed no correlation between the number of variants and gene length per base pair in each gene (*p* < 0.5). **(C)** Distribution of variants across the genomes of sequenced isolates. Notably, a cluster of genes that were identical in both groups had significantly high numbers of variants. Genes harboring more than 80 variants are indicated with their names.

Indeed, extensive analysis about the genome comparison between H99 and the clinical isolates identified polymorphisms in these two groups. **Fig 5A** showed variants that were common or specific in the isolates from HIV-u and HIV-i groups. We identified 55,292 common SNPs (we defined common variants as the ones that are found in more than 7 isolates) in both groups, and only 628 and 648 variants specific for HIV-u and HIV-i groups, respectively. Moreover, we found that compared to those specific SNPs of HIV-i group, the isolates of HIV-u group exhibited a significant higher level of variant enrichment at the intergenic region (54.05% vs 41.90%) but much less abundance in both upstream and exonic regions (4.98% vs 9.49% and 19.16% vs 28.18%, respectively). Other locations, such as intronic, downstream and ncRNA_exonic regions, showed no difference in variant distribution between the two groups **(Fig 5B)**. Furthermore, the 648 unique variants of isolates in HIV-u group were associated with 329 genes whereas the 628 unique variants in HIV-i group counts for 206 genes.

**Fig 5.**
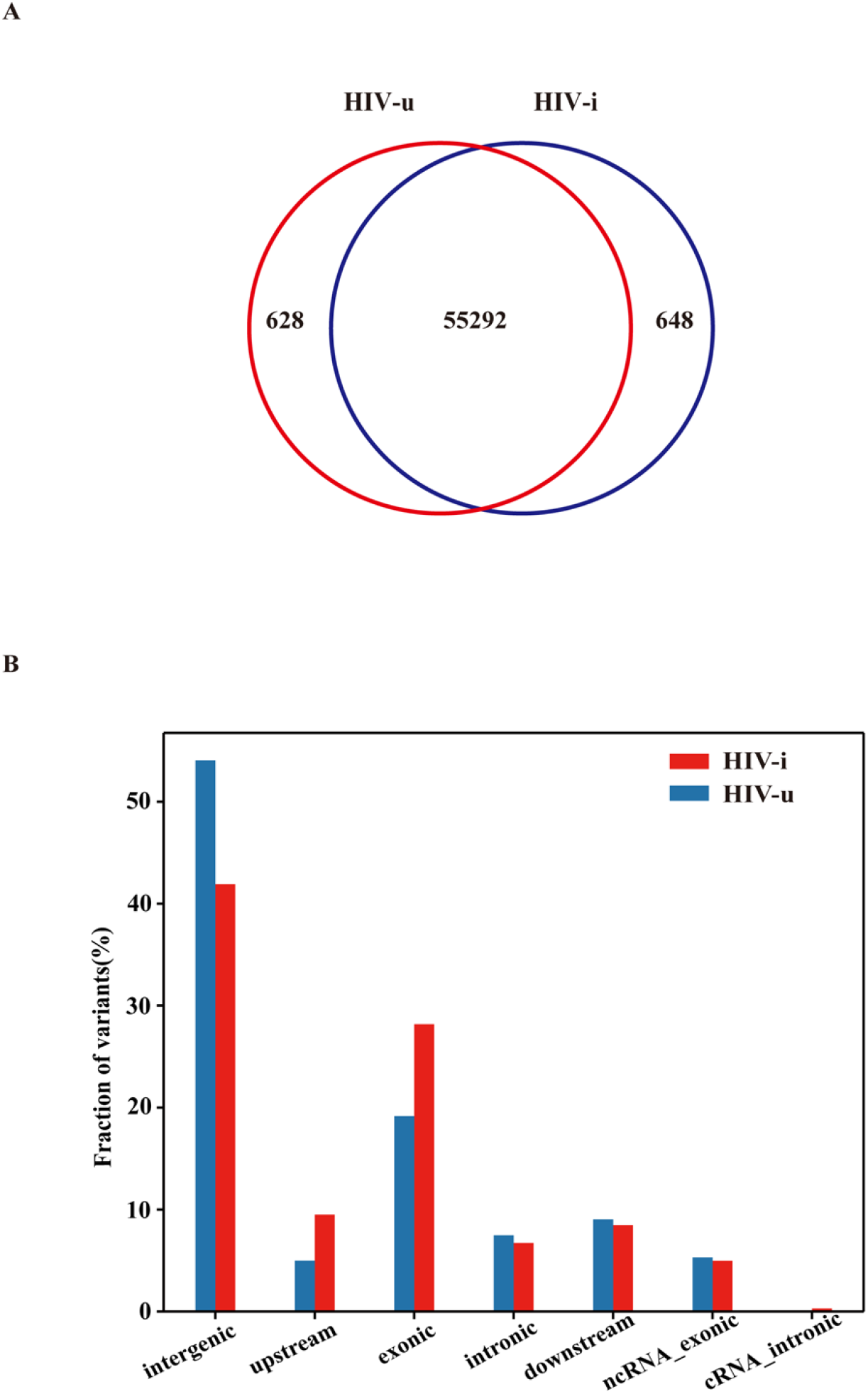
Common and group-specific variants identified in the sequenced isolates. **(A)** Venn diagram showing a summary of the number of common or group-specific variants. A common variant was defined as the one that are found in more than 7 isolates of each group. **(B)** Genomic distribution of the group-specific variants.

Importantly, when we determined the functional categories of the unique genes identified in each of the two groups using Gene Ontology (GO) Database, surprising findings were obtained. Overall, the functions of the specific genes of each group could be classified into three general directions (**Fig 6A, B** and **C**). In the biological process category, the top highly enriched GO terms in HIV-i group include protein glycosylation activity while those in HIV-u group were enriched in signal transduction activity. In the cellular component, the GO terms of RISC complex and riboflavin synthase complex were significantly enriched in HIV-i and HIV-u groups, respectively. As for the molecular function category, genes with specific variations in strains of HIV-i group showed a high percentage of metal/zinc ion binding process whereas those in HIV-u group dominated in the process of oxidoreductase activity. Although isolates from both HIV-i and HIV-u groups exhibit a large number of overlapping variants in the genomes, there is a subset of genes in each group that harbor unique mutations and account for different functions. The related functions and names of major mutated genes differed in two groups were listed in **Table 2**. We observed that the functions of HIV-i strain specific genes are mainly linked to microbial metabolism while those in HIV-u group impinge their roles in stress response, signal transduction and drug resistance. For example, the isolates from the HIV-u group harbored specific genetic variants in the genes *SSK1*, *TCO2* and *PBS2* who have been found to be key players in the fungal two-component system and the HOG signaling pathway and reported to regulate stress responses, drug sensitivity, sexual development, differentiation and virulence of *C. neoformans* [64, 65]. Moreover, specific mutations were also identified in the genes like *RHO104/RIM20* and *APT4* in the isolates from HIV-uninfected individuals but not in those from HIV-infected patients. Rho104/Rim20 are effectors of the PKA signaling pathway and required for many virulence-associated phenotypes such as titan cell formation in *C. neoformans* [66], and Apt4 is the P4-ATPase subunit of the Cdc50 family and regulates iron acquisition and virulence in *C. neoformans* [67]. In comparison, most genes harboring specific mutations in the isolates of HIV-i group are involved in metabolic processes, such as the MAPK signaling pathway (*SSK2*), tryptophan biosynthesis (*TRP1*), riboflavin biosynthesis (*RIB3*), cell wall protein glycosylation (*KTR3*) and cellular metabolism and compound biosynthesis (*KIC*, *HRK1*, *KIN1*, *GPA3*, *FZC45*, *SNF102*, *URE7*, *DST1* and *BCK1*).

**Table 2.** Characterization of major genes harboring specific variations in HIV-u and HIV-i groups, respectively

| Function | Broad annotation | Gene name | Description | Classic fisher |
| --- | --- | --- | --- | --- |
| **Genes harboring specific variations in HIV-u group** |  |  |  |  |
| Response to stress | CNAG_03818 | SSK1 | Response to fungicide/UV/radiation/light stimulus | 0.011 |
|  | CNAG_00769 | PBS2 | Cellular response to stimulus | 0.013 |
| Signaling | CNAG_05590 | TCO2 | Signaling, phosphorelay signal transduction system | 0.001 |
|  | CNAG_06606 | RHO104 | Intracellular signal transduction | 0.046 |
|  | CNAG_03751 | Unknown | Cation(calcium) channel activity | 0.022 |
|  | CNAG_02796 | Unknown | Transferase activity, transferring alkyl or aryl groups | 0.022 |
|  | CNAG_04599 | Unknown | 3-methyl-2-oxobutanoate hydroxymethy | 0.022 |
|  | CNAG_05712 | RIB4 | Intramolecular transferase activity, transferring acyl groups | 0.022 |
|  | CNAG_06806 | ETF1alpha |  |  |
| Antifungal resistance | CNAG_03582 | RIM20 | Macromolecule localization | 0.023 |
|  | CNAG_05282 | APT4 |  |  |
|  | CNAG_05395 | VAM6 |  |  |
|  | CNAG_02565 | Unknown | Organic acid biosynthetic process<br>Carboxylic acid biosynthetic process | 0.006 |
|  | CNAG_02796 | Unknown |  |  |
|  | CNAG_05194 | Unknown |  |  |
|  | CNAG_06421 | Unknown |  |  |
|  | CNAG_04087 | Unknown | Intrinsic and integral component of organelle membrane | 0.032 |
|  | CNAG_04556 | Unknown |  |  |
|  | CNAG_04604 | Unknown | Tryptophan-tRNA ligase activity | 0.022 |
|  | CNAG_06421 | Unknown | Ccetolactate synthase activity | 0.044 |
| Autophagy | CNAG_02796 | Unknown | Cellular amino acid metabolic and biosynthetic process | 0.020 |
|  | CNAG_04599 | Unknown |  |  |
|  | CNAG_06421 | Unknown |  |  |
|  | CNAG_04484 | Unknown | Organelle membrane fusion | 0.022 |
| **Genes harboring specific variations in HIV-i group** |  |  |  |  |
| MAPK | CNAG_05063 | SSK2 | MAPK/ protein serine/threonine/tyrosine kinase activity | 0.036 |
|  | CNAG_02853 | Unknown | Amidophosphoribosyltransferase activity | 0.036 |
| PHO pathway | CNAG_04501 | TRP1 | Indole-3-glycerol-phosphate synthase activity,<br>Phosphoribosylanthranilate isomerase activity | 0.036 |
|  | CNAG_02506 | RIB3 | 3,4-dihydroxy-2-butanone-4-phosphate synthase activity | 0.0361 |
| Glycosylation | CNAG_03832 | KTR3 | Glycoprotein metabolic and biosynthetic process, macromolecule glycosylation | 0.011 |
| Cellular metabolism and biosynthesis | CNAG_00405 | KIC1 | Regulation of cellular and biosynthetic metabolic process | 0.128 |
|  | CNAG_00745 | HRK1 | Regulation of biosynthetic process | 0.015 |
|  | CNAG_01938 | KIN1 | Biosynthetic process | 0.015 |
|  | CNAG_02090 | GPA3 | Carbohydrate derivative metabolic process | 0.012 |
|  | CNAG_02305 | FZC45 | Organic substance biosynthetic process | 0.006 |
|  | CNAG_06490 | SNF102 | Organic cyclic compound biosynthetic process | 0.007 |
|  | CNAG_00678 | URE7 | Regulation of biological process | 0.010 |
|  | CNAG_07679 | DST1 | Positive regulation of metabolic process | 0.015 |
|  | CNAG_04755 | BCK1 | Aromatic compound biosynthetic process | 0.019 |

**Fig 6.**
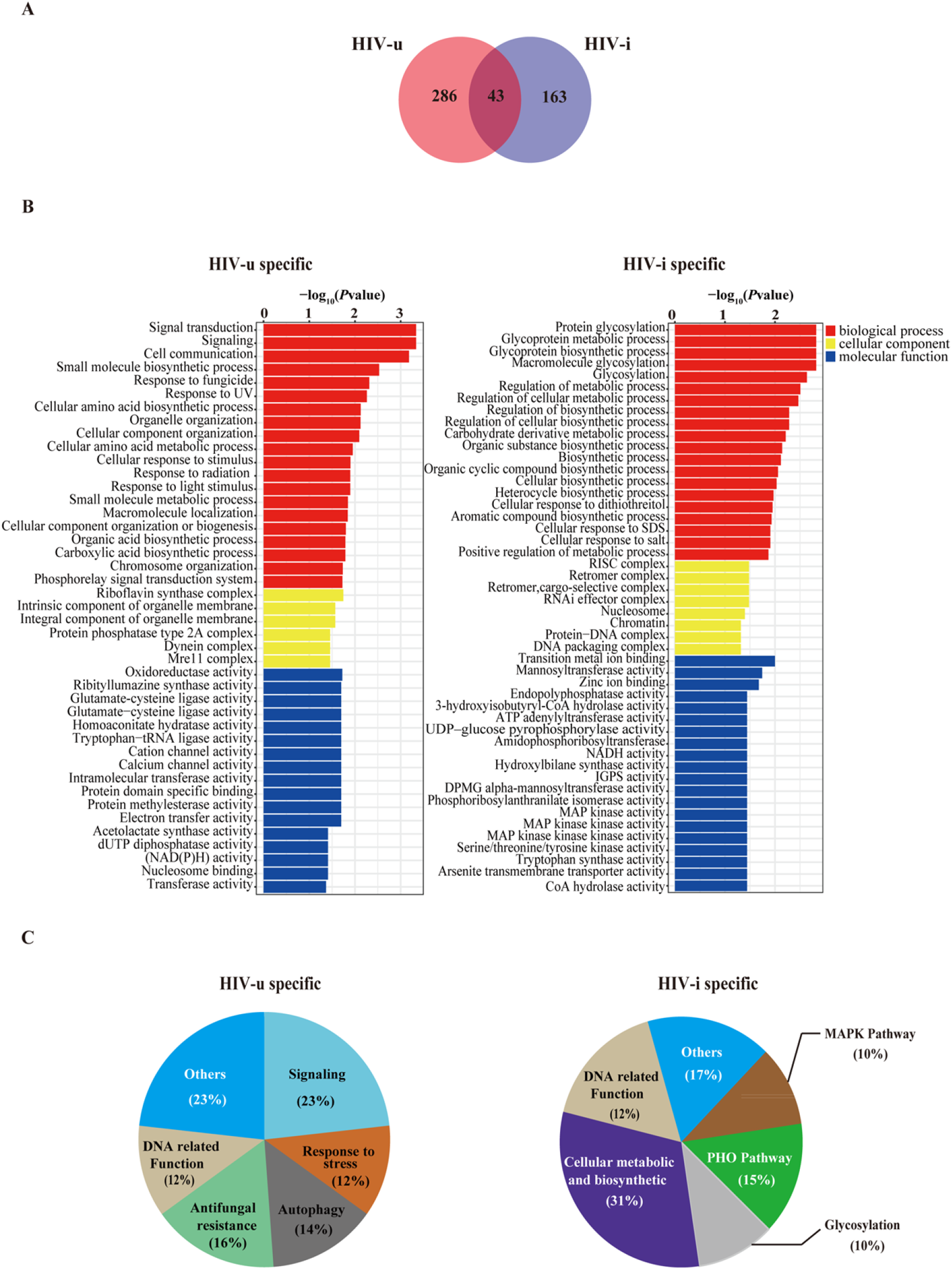
Characterization of the genes harboring group-specific variants. **(A)** Venn diagram showing the number of genes that were associated with group-specific variants. It has to be noted that 43 genes are common in both groups only because the variants in each group were mapped to different positions of the same gene. **(B)** Significantly enriched Gene Ontology (GO) categories (*p*-value <0.05) with group-specific genes. Functional classification pie chart showing the annotated genes that harbor group-specific variants, according to the Gene Ontology (GO) Term analysis

These data strongly suggest that compared to those isolated from HIV-infected patients, C. *neoformans* strains from HIV-uninfected individuals need to evolve genetic variations in a list of specific genes related to environmental adaptation, since microbes have to confront with more rigor host environment, such as immune clearance.

## Discussion

Cryptococcosis, primarily caused by *Cryptococcus neoformans* and *Cryptococcus gattii*, is a primary opportunistic fungal infection that has been found to be associated with patients with HIV infection. Of course, this fungal infection also occurs in other underlying disorders, including immunosuppressant usage, transplantation, cancers and diabetes mellitus *etc*. So far, studies on cryptococcosis in China have been carried out mainly in HIV-uninfected patients [29]. In this study, we used a large number of *C. neoformans* strains isolated from different sources, including those from the HIV-infected and HIV-uninfected patients, as well as natural environment, and systematically evaluated impacts of genetic and environmental factors on genotypes and virulence-related phenotypes of these isolates. We observed strong correlations and trends among the three different environments, which suggest that host environments appear to play important roles in affecting the virulence of *C. neoformans* isolates. Overall, strains isolated from the HIV-infected patients showed much lower virulence when compared to those from the HIV-uninfected individuals and a natural environment, indicating that *C. neoformans* pathogenicity has a great plasticity during interaction with different hosts. Interestingly, the information from whole-genome sequencing suggests that the high incidence of hotspot mutations may not be the major factor that accounts for the differential virulence among the isolates derived from either the HIV-i or HIV-u group. Instead, genomic variations within genes related to specific functions may act as a major driving force of host intrinsic virulence evolution.

Broad epidemiology and molecular typing studies of *C. neoformans* and *C. gattii* species complex have been reported all over the world, showing that genotypes are distinctly related to geographical locations [2]. Originally we planned to study the genotype-phenotype correlation using strains harboring different sequence types (ST). Unexpectedly, our results identified that all of the 105 isolates belong to the molecular type AFLP1/VNI and ST5 was the most common sequence type (n = 94; 89.5%), although other STs were also identified using MLST, making it impossible to evaluate the difference in the production of virulence factors between isolates from different STs. Consistently, the observation that ST5 was the predominant sequence type that caused human cryptococcosis in China was found to be the same case in most other Asian countries [68]. A distinct genotypic characteristic of Chinese *C. neoformans* isolates was apparent that ST5 represents the most prevalent genotype in clinical isolates from both HIV-infected and -uninfected samples and comparatively, the genotypes of those natural isolates (from pigeon droppings) are much more diverse. Furthermore, the data currently available imply that the genotypic diversity in isolates from HIV-infected patients appear to be more limited than those from HIV-uninfected individuals, and of course, supporting this notion requires larger isolate collections and decent work in data analyses.

Although strains with different genotypes harbor almost the same mating type, we observed differences of these strains in melanin production and capsule formation. Apparently, strains isolated from the HIV-uninfected patients produced significantly more melanin than those isolated from the HIV-infected patients and the nature, suggesting that once infected, a higher melanin biosynthesis of cryptococcal strains might be triggered in response to a relatively stronger immune system, given that host immune system has completely been dampened in HIV-infected patients. Considering the prevalence of L-DOPA in the central nervous system [27], our results also proposed that an increased melanin production in strains isolated from the HIV-uninfected patients could potentially occur *in vivo* and promotes virulence, leading to an increase in the mortality. Moreover, capsule generation in different strains yields similar results. A previous study showed that capsule size in cryptococcal cells was found to differ significantly between species and individual situation [69]. As expected, we observed the smallest size of capsule in strains from HIV-infected patients, more likely reflecting the fact that these strains might be more easily to cross biological barriers and to disseminate in the brain in immunocompromised patients. However, the relationships between capsule size and virulence have found controversial issues. One literature reported that *C. neoformans* isolates with higher capsule formation were associated with lower fungal clearance rates and increased intracranial pressure [70]. However, this notion was disagreed by other groups, as their studies showed that *C. neoformans* isolates with less capsule were more virulent and resulted in a higher fungal load in the brain [54]. In order to verify that *in vitro* capsule and melanin production does reflect the degree of pathogenicity, we test more virulence-related traits, including the ability of *C. neoformans* to survive in human CSF, the level of macrophage uptake, and morbidity in mouse model of cryptococcosis. These *in vitro*, *ex vivo* and *in vivo* data sustain the importance of host environment on the virulence evolution of *C. neoformans*. Actually, our findings are well consistent with the study by Robertson *et al*. [70]. Considering an impaired immune system in the HIV-infected patients, it is reasonable to conclude that virulence may not be an important factor in establishment of a successful infection. However, for colonization in a relatively healthy individual, such as HIV-negative patients, the cryptococcal strains must evolve to enhance their traits in virulence, in order to fight against the host immune system.

Mutation represents the ultimate source of the genetic variation required for adaptation and most of time the genomic mutation rate is adjusted to a level that best promote adaptation. We hypothesized that changes due to highly frequent genetic variations may contribute to the observed virulence variabilities among strains derived from different groups. To our surprise, we found that it is not the case, the basis of virulence attenuation and the multiple phenotypic changes of strains from HIV-infected group was unable to be largely addressed by the genomic changes at a high frequency. Consistently, a previous whole-genome sequencing study in 32 isolates from 18 South African patients with recurrent cryptococcal meningitis also revealed that only a few genetic changes, including single nucleotide polymorphism (SNPs), small insertions/deletions (indels) and variation in copy number, were identified between incident and replapse isolates [71]. However, we did observe genetic variations for a few specific genes in these two groups. For example, the isolates from HIV-uninfected group harbor frequent genomic mutations identified in a list of genes specifically related to environmental adaptation, which could explain the increased virulent phenotypes of this group when compared to the strains from HIV-infected group. Genetic mutations actively respond to the change of host environment and thereby drive the adaptive evolution of fungal virulence. In contrast, we observed in HIV-infected group that genetic variations were mostly occurred in genes associated with metabolic processes, implying that the mutations may significantly correlate with survival with respect to the possible role of these genes in receiving nutrients from the host, which is expected to have almost no immune stresses.

In conclusion, through successful combination of genotyping, pathogenic phenotyping and comparative genomics, our work has for the first time performed phenotypic characterization and evaluated virulence-related properties of *C. neoformans* strains isolated from three different host environments, including the HIV-uninfected individuals, HIV-infected patients and the natural resources. Our studies highlighted that host environmental factors may majorly account for the attenuation of virulence and various other phenotypic changes in the strains from HIV-infected patients that are different from their HIV-uninfected counterparts. Moreover, our work also supports an essential role of the genetic variations in driving the evolution of fungal virulence. This might be extremely important for *C. neoformans* to avoid clearance of host immunity and establish successful colonization. Further biochemical, molecular genetics and immunological studies are required to confirm and assess the relative importance of both environmental and genetic factors in regulating pathogenicity and causing invasive infection of *C. neoformans*. Such studies will not only improve our understanding of pathogenic mechanisms of this fungus but also facilitate design of new therapeutic approaches based on these factors.

## Acknowledgments

The authors would like to convey gratitude to Prof. Liao Wanqing (Department of Dermatology, Changzheng Hospital, Second Military University, Shanghai, China), Prof. Xue Chaoyang (The Rutgers University, USA) for valuable comments, sharing strains and providing human CSF samples. The authors also thank all members of the Chen-lab for helpful advice and discussion. CC is supported by grants from the MOST Key R&D Program (2020YFA0907200); National Natural Science Foundation of China (31870141, 31570140); the Key Research Program of the Chinese Academy of Sciences (KGFZD-135-19-11,153831KYSB20170043); the Innovation Capacity Building Project of Jiangsu Province (BM2020019). JZ is supported by the National Natural Science Foundation of China (81874240) and Natural Science Foundation of Shanghai (16ZR1432400). HL is supported by the Special Program (19SWAQ18) and Open Project of the State Key Laboratory of Trauma, Burns and Combined Injury (SKLKF201803). XH is supported by the National Natural Science Foundation of China to (32070146, 31600119); Youth Innovation Promotion Association, CAS; Natural Science Foundation of Shanghai (20ZR1463800, 15ZR1444400). CC and JZ conceived and designed the research study and wrote the paper; WY and YY performed the experiments, analyzed the data and wrote the paper; LL and GL collected the strains from HIV-infected patients and analyzed the data; XC, ML, TZ, HL, XH and TJ analyzed the data. LZ and MC provided strains from HIV-uninfected patients and the natural resources. All authors read and approved the final version of the paper.

## Conflict of interest

The authors declare that they have no conflict of interest.

## Notes

### Competing Interest Statement

The authors have declared no competing interest.

## References

1. Kwon-Chung KJ, Varma A. Do major species concepts support one, two or more species within Cryptococcus neoformans? FEMS Yeast Res. 2006;6(4):574–87. doi: 10.1111/j.1567-1364.2006.00088.x. PubMed PMID: 16696653.

2. Cogliati M. Global Molecular Epidemiology of Cryptococcus neoformans and Cryptococcus gattii: An Atlas of the Molecular Types. Scientifica (Cairo). 2013;2013:675213. doi: 10.1155/2013/675213. PubMed PMID: 24278784; PubMed Central PMCID: PMCPMC3820360.

3. Kidd SE, Hagen F, Tscharke RL, Huynh M, Bartlett KH, Fyfe M, et al. A rare genotype of Cryptococcus gattii caused the cryptococcosis outbreak on Vancouver Island (British Columbia, Canada). Proc Natl Acad Sci U S A. 2004;101(49):17258–63. doi: 10.1073/pnas.0402981101. PubMed PMID: 15572442; PubMed Central PMCID: PMCPMC535360.

4. Guo LY, Liu LL, Liu Y, Chen TM, Li SY, Yang YH, et al. Characteristics and outcomes of cryptococcal meningitis in HIV seronegative children in Beijing, China, 2002-2013. BMC Infect Dis. 2016;16(1):635. doi: 10.1186/s12879-016-1964-6. PubMed PMID: 27814690; PubMed Central PMCID: PMCPMC5097362.

5. Chen M, Xu Y, Hong N, Yang Y, Lei W, Du L, et al. Epidemiology of fungal infections in China. Front Med. 2018;12(1):58–75. doi: 10.1007/s11684-017-0601-0. PubMed PMID: 29380297.

6. Rajasingham R, Smith RM, Park BJ, Jarvis JN, Govender NP, Chiller TM, et al. Global burden of disease of HIV-associated cryptococcal meningitis: an updated analysis. Lancet Infect Dis. 2017;17(8):873–81. doi: 10.1016/S1473-3099(17)30243-8. PubMed PMID: 28483415; PubMed Central PMCID: PMCPMC5818156.

7. Wu SY, Lei Y, Kang M, Xiao YL, Chen ZX. Molecular characterisation of clinical Cryptococcus neoformans and Cryptococcus gattii isolates from Sichuan province, China. Mycoses. 2015;58(5):280–7. doi: 10.1111/myc.12312. PubMed PMID: 25808662.

8. Alves GS, Freire AK, Bentes Ados S, Pinheiro JF, de Souza JV, Wanke B, et al. Molecular typing of environmental Cryptococcus neoformans/C. gattii species complex isolates from Manaus, Amazonas, Brazil. Mycoses. 2016;59(8):509–15. doi: 10.1111/myc.12499. PubMed PMID: 27005969.

9. Benham RW. Cryptococcosis and blastomycosis. Ann N Y Acad Sci. 1950;50(10):1299–314. doi: 10.1111/j.1749-6632.1950.tb39828.x. PubMed PMID: 14783320.

10. Evans EE. The antigenic composition of Cryptococcus neoformans. I. A serologic classification by means of the capsular and agglutination reactions. J Immunol. 1950;64(5):423–30. PubMed PMID: 15415610.

11. Wilson DE, Bennett JE, Bailey JW. Serologic grouping of Cryptococcus neoformans. Proc Soc Exp Biol Med. 1968;127(3):820–3. doi: 10.3181/00379727-127-32812. PubMed PMID: 5651140.

12. Kwon-Chung KJ, Boekhout T, Fell JW, Diaz M. (1557) Proposal to conserve the name Cryptococcus gattii against C. hondurianus and C. bacillisporus (Basidiomycota, Hymenomycetes, Tremellomycetidae). Taxon. 2002;51(4):804–6.

13. Park BJ, Wannemuehler KA, Marston BJ, Govender N, Pappas PG, Chiller TM. Estimation of the current global burden of cryptococcal meningitis among persons living with HIV/AIDS. Aids. 2009;23(4):525–30.

14. Dromer F, Mathoulin S, Dupont B, Letenneur L, Ronin O, Group FCS. Individual and environmental factors associated with infection due to Cryptococcus neoformans serotype D. Clinical infectious diseases. 1996;23(1):91–6.

15. Kwon-Chung KJ, Edman JC, Wickes BL. Genetic association of mating types and virulence in Cryptococcus neoformans. Infection and immunity. 1992;60(2):602–5.

16. Meyer W, Castaneda A, Jackson S, Huynh M, Castaneda E, IberoAmerican Cryptococcal Study G. Molecular typing of IberoAmerican Cryptococcus neoformans isolates. Emerg Infect Dis. 2003;9(2):189–95. doi: 10.3201/eid0902.020246. PubMed PMID: 12603989; PubMed Central PMCID: PMCPMC2901947.

17. Boekhout T, Theelen B, Diaz M, Fell JW, Hop WCJ, Abeln ECA, et al. Hybrid genotypes in the pathogenic yeast Cryptococcus neoformans. Microbiology (Reading). 2001;147(Pt 4):891–907. doi: 10.1099/00221287-147-4-891. PubMed PMID: 11283285.

18. Meyer W, Aanensen DM, Boekhout T, Cogliati M, Diaz MR, Esposto MC, et al. Consensus multi-locus sequence typing scheme for Cryptococcus neoformans and Cryptococcus gattii. Med Mycol. 2009;47(6):561–70. doi: 10.1080/13693780902953886. PubMed PMID: 19462334; PubMed Central PMCID: PMCPMC2884100.

19. Cuomo CA, Rhodes J, Desjardins CA. Advances in Cryptococcus genomics: insights into the evolution of pathogenesis. Mem Inst Oswaldo Cruz. 2018;113(7):e170473. doi: 10.1590/0074-02760170473. PubMed PMID: 29513784; PubMed Central PMCID: PMCPMC5851040.

20. Hagen F, Colom MF, Swinne D, Tintelnot K, Iatta R, Montagna MT, et al. Autochthonous and dormant Cryptococcus gattii infections in Europe. Emerg Infect Dis. 2012;18(10):1618–24. doi: 10.3201/eid1810.120068. PubMed PMID: 23017442; PubMed Central PMCID: PMCPMC3471617.

21. Antinori S. New Insights into HIV/AIDS-Associated Cryptococcosis. ISRN AIDS. 2013;2013:471363. doi: 10.1155/2013/471363. PubMed PMID: 24052889; PubMed Central PMCID: PMCPMC3767198.

22. Litvintseva AP, Thakur R, Vilgalys R, Mitchell TG. Multilocus sequence typing reveals three genetic subpopulations of Cryptococcus neoformans var. grubii (serotype A), including a unique population in Botswana. Genetics. 2006;172(4):2223–38. doi: 10.1534/genetics.105.046672. PubMed PMID: 16322524; PubMed Central PMCID: PMCPMC1456387.

23. Andrade-Silva LE, Ferreira-Paim K, Ferreira TB, Vilas-Boas A, Mora DJ, Manzato VM, et al. Genotypic analysis of clinical and environmental Cryptococcus neoformans isolates from Brazil reveals the presence of VNB isolates and a correlation with biological factors. PLoS One. 2018;13(3):e0193237. doi: 10.1371/journal.pone.0193237. PubMed PMID: 29505557; PubMed Central PMCID: PMCPMC5837091.

24. Viviani MA, Cogliati M, Esposto MC, Lemmer K, Tintelnot K, Colom Valiente MF, et al. Molecular analysis of 311 Cryptococcus neoformans isolates from a 30-month ECMM survey of cryptococcosis in Europe. FEMS Yeast Res. 2006;6(4):614–9. doi: 10.1111/j.1567-1364.2006.00081.x. PubMed PMID: 16696657.

25. Altamirano S, Jackson KM, Nielsen K. The interplay of phenotype and genotype in Cryptococcus neoformans disease. Biosci Rep. 2020;40(10). doi: 10.1042/BSR20190337. PubMed PMID: 33021310; PubMed Central PMCID: PMCPMC7569153.

26. Perfect JR, Dismukes WE, Dromer F, Goldman DL, Graybill JR, Hamill RJ, et al. Clinical practice guidelines for the management of cryptococcal disease: 2010 update by the infectious diseases society of america. Clin Infect Dis. 2010;50(3):291–322. doi: 10.1086/649858. PubMed PMID: 20047480; PubMed Central PMCID: PMCPMC5826644.

27. Wiesner DL, Moskalenko O, Corcoran JM, McDonald T, Rolfes MA, Meya DB, et al. Cryptococcal genotype influences immunologic response and human clinical outcome after meningitis. mBio. 2012;3(5). doi: 10.1128/mBio.00196-12. PubMed PMID: 23015735; PubMed Central PMCID: PMCPMC3448160.

28. Beale MA, Sabiiti W, Robertson EJ, Fuentes-Cabrejo KM, O’Hanlon SJ, Jarvis JN, et al. Genotypic Diversity Is Associated with Clinical Outcome and Phenotype in Cryptococcal Meningitis across Southern Africa. PLoS Negl Trop Dis. 2015;9(6):e0003847. doi: 10.1371/journal.pntd.0003847. PubMed PMID: 26110902; PubMed Central PMCID: PMCPMC4482434.

29. Chen J, Varma A, Diaz MR, Litvintseva AP, Wollenberg KK, Kwon-Chung KJ. Cryptococcus neoformans strains and infection in apparently immunocompetent patients, China. Emerg Infect Dis. 2008;14(5):755–62. doi: 10.3201/eid1405.071312. PubMed PMID: 18439357; PubMed Central PMCID: PMCPMC2600263.

30. Khayhan K, Hagen F, Pan W, Simwami S, Fisher MC, Wahyuningsih R, et al. Geographically structured populations of Cryptococcus neoformans Variety grubii in Asia correlate with HIV status and show a clonal population structure. PLoS One. 2013;8(9):e72222. doi: 10.1371/journal.pone.0072222. PubMed PMID: 24019866; PubMed Central PMCID: PMCPMC3760895.

31. Day JN, Qihui S, Thanh LT, Trieu PH, Van AD, Thu NH, et al. Comparative genomics of Cryptococcus neoformans var. grubii associated with meningitis in HIV infected and uninfected patients in Vietnam. PLoS Negl Trop Dis. 2017;11(6):e0005628. doi: 10.1371/journal.pntd.0005628. PubMed PMID: 28614360; PubMed Central PMCID: PMCPMC5484541.

32. Dou HT, Xu YC, Wang HZ, Li TS. Molecular epidemiology of Cryptococcus neoformans and Cryptococcus gattii in China between 2007 and 2013 using multilocus sequence typing and the DiversiLab system. Eur J Clin Microbiol Infect Dis. 2015;34(4):753–62. doi: 10.1007/s10096-014-2289-2. PubMed PMID: 25471194.

33. Simwami SP, Khayhan K, Henk DA, Aanensen DM, Boekhout T, Hagen F, et al. Low diversity Cryptococcus neoformans variety grubii multilocus sequence types from Thailand are consistent with an ancestral African origin. PLoS Pathog. 2011;7(4):e1001343. doi: 10.1371/journal.ppat.1001343. PubMed PMID: 21573144; PubMed Central PMCID: PMCPMC3089418.

34. Perfect JR. Cryptococcus neoformans: a sugar-coated killer with designer genes. FEMS Immunol Med Microbiol. 2005;45(3):395–404. doi: 10.1016/j.femsim.2005.06.005. PubMed PMID: 16055314.

35. Beenhouwer DO, Shapiro S, Feldmesser M, Casadevall A, Scharff MD. Both Th1 and Th2 cytokines affect the ability of monoclonal antibodies to protect mice against Cryptococcus neoformans. Infect Immun. 2001;69(10):6445–55. doi: 10.1128/IAI.69.10.6445-6455.2001. PubMed PMID: 11553589; PubMed Central PMCID: PMCPMC98780.

36. Kozel TR. Virulence factors of Cryptococcus neoformans. Trends Microbiol. 1995;3(8):295–9. PubMed PMID: 8528612.

37. Zaragoza O. Basic principles of the virulence of Cryptococcus. Virulence. 2019;10(1):490–501. doi: 10.1080/21505594.2019.1614383. PubMed PMID: 31119976; PubMed Central PMCID: PMCPMC6550552.

38. Mayer FL, Kronstad JW. Disarming Fungal Pathogens: Bacillus safensis Inhibits Virulence Factor Production and Biofilm Formation by Cryptococcus neoformans and Candida albicans. MBio. 2017;8(5). doi: 10.1128/mBio.01537-17. PubMed PMID: 28974618; PubMed Central PMCID: PMCPMC5626971.

39. Thompson JD, Higgins DG, Gibson TJ. CLUSTAL W: improving the sensitivity of progressive multiple sequence alignment through sequence weighting, position-specific gap penalties and weight matrix choice. Nucleic Acids Res. 1994;22(22):4673–80. PubMed PMID: 7984417; PubMed Central PMCID: PMCPMC308517.

40. Liu OW, Chun CD, Chow ED, Chen C, Madhani HD, Noble SM. Systematic genetic analysis of virulence in the human fungal pathogen Cryptococcus neoformans. Cell. 2008;135(1):174–88. doi: 10.1016/j.cell.2008.07.046. PubMed PMID: 18854164; PubMed Central PMCID: PMCPMC2628477.

41. García-Rodas R, Cordero RJ, Trevijano-Contador N, Janbon G, Moyrand F, Casadevall A, et al. Capsule growth in Cryptococcus neoformans is coordinated with cell cycle progression. MBio. 2014;5(3):e00945–14.

42. Fernandes OdFL, Costa CR, Junior RdSL, Vinaud MC, e Souza LKH, de Paula JAM, et al. Effects of Pimenta pseudocaryophyllus (Gomes) LR Landrum, on melanized and non-melanized Cryptococcus neoformans. Mycopathologia. 2012;174(5-6):421–8.

43. Shulman JJ, Hontz A, Sedlis A, Walters AT, Balin H, LoSCUITO L. The Pap smear: take two. American journal of obstetrics and gynecology. 1975;121(8):1024–8.

44. Hansakon A, Mutthakalin P, Ngamskulrungroj P, Chayakulkeeree M, Angkasekwinai P. Cryptococcus neoformans and Cryptococcus gattii clinical isolates from Thailand display diverse phenotypic interactions with macrophages. Virulence. 2019;10(1):26–36.

45. Sabiiti W, Robertson E, Beale MA, Johnston SA, Brouwer AE, Loyse A, et al. Efficient phagocytosis and laccase activity affect the outcome of HIV-associated cryptococcosis. The Journal of clinical investigation. 2014;124(5):2000–8.

46. Denham ST, Verma S, Reynolds RC, Worne CL, Daugherty JM, Lane TE, et al. Regulated release of cryptococcal polysaccharide drives virulence and suppresses immune infiltration into the central nervous system. Infect Immun. 2017. doi: 10.1128/IAI.00662-17. PubMed PMID: 29203547; PubMed Central PMCID: PMCPMC5820953.

47. Guess T, Lai H, Smith SE, Sircy L, Cunningham K, Nelson DE, et al. Size Matters: Measurement of Capsule Diameter in Cryptococcus neoformans. J Vis Exp. 2018;(132). doi: 10.3791/57171. PubMed PMID: 29553511.

48. Tugume L, Rhein J, Hullsiek KH, Mpoza E, Kiggundu R, Ssebambulidde K, et al. HIV-associated cryptococcal meningitis occurring at relatively higher CD4 counts. The Journal of infectious diseases. 2019;219(6):877–83.

49. Tamura K, Stecher G, Peterson D, Filipski A, Kumar S. MEGA6: Molecular Evolutionary Genetics Analysis version 6.0. Mol Biol Evol. 2013;30(12):2725–9. doi: 10.1093/molbev/mst197. PubMed PMID: 24132122; PubMed Central PMCID: PMCPMC3840312.

50. Dou H, Wang H, Xie S, Chen X, Xu Z, Xu Y. Molecular characterization of Cryptococcus neoformans isolated from the environment in Beijing, China. Med Mycol. 2017;55(7):737–47. doi: 10.1093/mmy/myx026. PubMed PMID: 28431114.

51. Hull CM, Heitman J. Genetics of Cryptococcus neoformans. Annu Rev Genet. 2002;36:557–615. doi: 10.1146/annurev.genet.36.052402.152652. PubMed PMID: 12429703.

52. Wickes BL. The role of mating type and morphology in Cryptococcus neoformans pathogenesis. Int J Med Microbiol. 2002;292(5-6):313–29. doi: 10.1078/1438-4221-00216. PubMed PMID: 12452279.

53. Nielsen K, Heitman J. Sex and virulence of human pathogenic fungi. Adv Genet. 2007;57:143–73. doi: 10.1016/S0065-2660(06)57004-X. PubMed PMID: 17352904.

54. Pool A, Lowder L, Wu Y, Forrester K, Rumbaugh J. Neurovirulence of Cryptococcus neoformans determined by time course of capsule accumulation and total volume of capsule in the brain. J Neurovirol. 2013;19(3):228–38. doi: 10.1007/s13365-013-0169-7. PubMed PMID: 23733307.

55. Eisenman HC, Chow SK, Tse KK, McClelland EE, Casadevall A. The effect of L-DOPA on Cryptococcus neoformans growth and gene expression. Virulence. 2011;2(4):329–36. doi: 10.4161/viru.2.4.16136. PubMed PMID: 21705857; PubMed Central PMCID: PMCPMC3173677.

56. Johnston SA, May RC. Cryptococcus interactions with macrophages: evasion and manipulation of the phagosome by a fungal pathogen. Cellular microbiology. 2013;15(3):403–11.

57. Smith LM, Dixon EF, May RC. The fungal pathogen C ryptococcus neoformans manipulates macrophage phagosome maturation. Cellular microbiology. 2015;17(5):702–13.

58. García-Rodas R, Zaragoza O. Catch me if you can: phagocytosis and killing avoidance by Cryptococcus neoformans. FEMS Immunology & Medical Microbiology. 2012;64(2):147–61.

59. Casadevall A, Coelho C, Cordero RJ, Dragotakes Q, Jung E, Vij R, et al. The capsule of Cryptococcus neoformans. Virulence. 2019;10(1):822–31.

60. Gish SR, Maier EJ, Haynes BC, Santiago-Tirado FH, Srikanta DL, Ma CZ, et al. Computational Analysis Reveals a Key Regulator of Cryptococcal Virulence and Determinant of Host Response. MBio. 2016;7(2):e00313–16. doi: 10.1128/mBio.00313-16. PubMed PMID: 27094327; PubMed Central PMCID: PMCPMC4850258.

61. Wang ZA, Li LX, Doering TL. Unraveling synthesis of the cryptococcal cell wall and capsule. Glycobiology. 2018;28(10):719–30. doi: 10.1093/glycob/cwy030. PubMed PMID: 29648596; PubMed Central PMCID: PMCPMC6142866.

62. Casadevall A, Pirofski LA. Host-pathogen interactions: redefining the basic concepts of virulence and pathogenicity. Infect Immun. 1999;67(8):3703–13. doi: 10.1128/IAI.67.8.3703-3713.1999. PubMed PMID: 10417127; PubMed Central PMCID: PMCPMC96643.

63. Urban M, Pant R, Raghunath A, Irvine AG, Pedro H, Hammond-Kosack KE. The Pathogen-Host Interactions database (PHI-base): additions and future developments. Nucleic Acids Res. 2015;43(Database issue):D645–55. doi: 10.1093/nar/gku1165. PubMed PMID: 25414340; PubMed Central PMCID: PMCPMC4383963.

64. Bahn YS, Kojima K, Cox GM, Heitman J. A unique fungal two-component system regulates stress responses, drug sensitivity, sexual development, and virulence of Cryptococcus neoformans. Mol Biol Cell. 2006;17(7):3122–35. doi: 10.1091/mbc.e06-02-0113. PubMed PMID: 16672377; PubMed Central PMCID: PMCPMC1483045.

65. Bahn YS, Kojima K, Cox GM, Heitman J. Specialization of the HOG pathway and its impact on differentiation and virulence of Cryptococcus neoformans. Mol Biol Cell. 2005;16(5):2285–300. doi: 10.1091/mbc.e04-11-0987. PubMed PMID: 15728721; PubMed Central PMCID: PMCPMC1087235.

66. Okagaki LH, Wang Y, Ballou ER, O’Meara TR, Bahn YS, Alspaugh JA, et al. Cryptococcal titan cell formation is regulated by G-protein signaling in response to multiple stimuli. Eukaryot Cell. 2011;10(10):1306–16. doi: 10.1128/EC.05179-11. PubMed PMID: 21821718; PubMed Central PMCID: PMCPMC3187071.

67. Hu G, Caza M, Bakkeren E, Kretschmer M, Bairwa G, Reiner E, et al. A P4-ATPase subunit of the Cdc50 family plays a role in iron acquisition and virulence in Cryptococcus neoformans. Cell Microbiol. 2017;19(6). doi: 10.1111/cmi.12718. PubMed PMID: 28061020; PubMed Central PMCID: PMCPMC5429215.

68. Choi YH, Ngamskulrungroj P, Varma A, Sionov E, Hwang SM, Carriconde F, et al. Prevalence of the VNIc genotype of Cryptococcus neoformans in non-HIV-associated cryptococcosis in the Republic of Korea. FEMS Yeast Res. 2010;10(6):769–78. doi: 10.1111/j.1567-1364.2010.00648.x. PubMed PMID: 20561059; PubMed Central PMCID: PMCPMC2920376.

69. Fernandes KE, Brockway A, Haverkamp M, Cuomo CA, van Ogtrop F, Perfect JR, et al. Phenotypic Variability Correlates with Clinical Outcome in Cryptococcus Isolates Obtained from Botswanan HIV/AIDS Patients. MBio. 2018;9(5). doi: 10.1128/mBio.02016-18. PubMed PMID: 30352938; PubMed Central PMCID: PMCPMC6199498.

70. Robertson EJ, Najjuka G, Rolfes MA, Akampurira A, Jain N, Anantharanjit J, et al. Cryptococcus neoformans ex vivo capsule size is associated with intracranial pressure and host immune response in HIV-associated cryptococcal meningitis. J Infect Dis. 2014;209(1):74–82. doi: 10.1093/infdis/jit435. PubMed PMID: 23945372; PubMed Central PMCID: PMCPMC3864387.

71. Chen Y, Farrer RA, Giamberardino C, Sakthikumar S, Jones A, Yang T, et al. Microevolution of Serial Clinical Isolates of Cryptococcus neoformans var. grubii and C. gattii. mBio. 2017;8(2). doi: 10.1128/mBio.00166-17. PubMed PMID: 28270580; PubMed Central PMCID: PMCPMC5340869.

